# Log-order improved *in trans* hammerhead ribozyme turnover rates: reevaluating therapeutic space for small catalytic RNAs

**DOI:** 10.1101/2022.04.26.489422

**Authors:** Jason M. Myers, Jack M. Sullivan

## Abstract

We discovered an enhanced functionality hammerhead ribozyme (**EhhRz**), designed to act *in trans* against human rod opsin (*RHO*) mRNA, with turnover activity >300 nM min^−1^ under substrate-excess conditions and physiological Mg^2+^ levels (1 mM). We developed a real-time moderate-throughput fluorescence quantitative hhRz kinetic assay, which is linear with substrate and product moles and supported by gel-based measures. The EhhRz targets a CUC↓ cleavage site in a substrate with no predicted secondary/tertiary structure and demonstrates classic Michaelis-Menten turnover behavior when the substrate is in 10-fold excess (*V*_max_/*K_m_* up to 1.60 × 10^8^ min^−1^ M^−1^), which is comparable to RNase A. EhhRzs show cooperative titration with a *K_d_* of 0.73 ± 0.02 mM at cellular Mg^2+^ concentrations and a Hill coefficient of 1.73 ± 0.07. The upstream EhhRz antisense flank (bound to a downstream substrate flank) interacts with stem-loop II, and examinations of different variants revealed that a U7 residue in the downstream flank of the substrate is *not* essential for enhanced activity. Under single-turnover conditions with substrate pre-annealed to enzyme, reaction rates exceeded 1,000 min^−1^. These findings show that RNA catalysis approaches the efficiency of the ribosome and suggests EhhRz *in trans* is a druggable nucleic acid therapeutic.

## INTRODUCTION

Small catalytic RNAs (class of noncoding RNAs) have been studied for more than 3 decades since the paradigm-shifting discoveries of Altman and Cech that RNA chemistry was capable of catalysis (Altman, 1990; Cech, 1990). The hammerhead ribozyme (hhRz) is a catalytic RNA with a highly conserved 11-nt core enzyme sequence (Uhlenbeck, 1987). The minimal hhRz (**mhhRz**) was isolated by Uhlenbeck from larger virusoid RNAs that function efficiently but variably in rolling circle RNA replication *in cis* (Uhlenbeck, 1987; Steponovskaya and Uhlenbeck, 2008). The mhhRz can function as a Michaelis-Menten enzyme under substrate-excess conditions with a minimal target RNA with no structure and non-physiological free Mg^2+^ levels (generally ≥ 10 mM). There was initial hope that the mhhRz could serve as an engineered site-specific RNA endonuclease for nucleic acid therapeutics (Usman et al., 1996; Birikh et al. 1997). However, mhhRzs exhibit slow catalysis, generally a 1–2- min^−1^ turnover for short targets pre-annealed with excess enzyme and often orders of magnitude lower for structured RNAs with constrained accessibility to annealing (Stage-Zimmermann and Uhlenbeck, 1998; Breaker et al., 2003; Emilsson et al., 2003). The latter can be at least partially mitigated by accessibility prediction algorithms and experimental target mRNA mapping approaches (Patzel et al., Abdelmaksoud et al. 2009; Yau et al., 2016; Froebel et al., 2018; Yau et al., 2019; Smola and Weeks, 2018; Busan et al., 2019). Because any therapeutic catalytic RNA will add a component to the intrinsic rate of degradation, only stable mRNAs (long half-lives) encoding structural or enzymatic proteins are viable targets. The extremely slow turnover rate at physiological intracellular Mg^2+^ levels is the major factor in therapeutic applications of mhhRzs because it necessitates high therapeutic enzyme [E] levels to achieve significant knockdown of the mRNA substrate [S], even at rare, fully accessible regions. These are conditions far from Michaelis-Menten kinetics (i.e., [S]>>[E]). The need for increased delivery or expression of the therapeutic results in not only greater technological challenges but also increased risks for off-target effects and vector- or therapeutic-related toxicity (Cideciyian et al., 2018).

The discovery of upstream tertiary accessory elements (TAEs) of native or extended hhRzs (**xhhRzs**) in 2003–2004 (Khvorova et al., 2003; De La Peña et al., 2003; Peneda et al., 2004), long missing from 15 years of study of the mhhRz (Uhlenbeck, 2003), has proven essential to understanding RNA catalytic function at cellular levels of Mg^2+^ (0.4–2 mM). xhhRzs tend to have improved catalytic rates when operating natively *in cis* (intramolecular), but the therapeutic *in trans* (intermolecular) configuration has limitations. The upstream antisense flank must be extended to insert the TAE, which could decrease specificity and suppress product leaving rates, which would constrain enzyme turnover capacity under steady-state conditions (Saksmerprome et al., 2004; Roychowdhury et al., 2006). The structure of xhhRzs revealed that the upstream TAE interacts with the loop capping stem-loop II of the enzyme core to stabilize a catalytically active conformation at a low Mg^2+^, which was only populated in the mhhRz at high Mg^2+^ (Blount and Uhlenbeck, 2005; Martick and Scott, 2006; Nelson and Uhlenbeck, 2006). Studies have since illustrated that there are critical residues in the highly conserved cores of the xhhRz and mhhRz, such as G12 and G8, which function as general base and acid drivers, respectively, for RNA phosphodiester cleavage, similarly to histidine residues in RNase A (Tang and Breaker, 1997; Nelson and Uhlenbeck, Thompson et al., 1995; Strobel and Cochrane, 2007; Cochrane and Strobel, 2008).

Because many naturally occurring xhhRzs have a U7 residue in the target substrate sequence 7 nt downstream of the cleavage site (Shepontinovskaya and Uhlenbeck, 2008; Dufour et al., 2009), the Scott group (O’Rourke et al., 2015) suggested that an unpaired substrate U7 residue forms a Hoogsteen face base interaction with the fourth A in the GUGA (or GUUA) tetraloop capping stem II loop. Single-turnover rates (enzyme excess under pre-annealed conditions) at 10 mM Mg^2+^ and 27°C were 61–95 min^−1^, with rates extrapolated mathematically from measurements at pH 5.6. Their study demonstrated that rate enhancements of hhRzs do not require complex structured upstream TAEs, but they do require certain *in trans* tertiary interactions. Unfortunately, functional assays were not evaluated at physiological [Mg^2+^].

In this study, we discovered that enhanced hhRzs (EhhRzs) *in trans*, without upstream TAEs, can achieve rates >370 nM min^−1^ under Michaelis-Menten ([S]>>[E]) conditions at physiological temperature and Mg^2+^ concentrations (≤1 mM). EhhRzs have catalytic turnover (*V*_max_/*K_m_*) on the same scale as the highly efficient protein enzyme RNase A. Moreover, we have substantially extended the findings of O’Rourke et al. (2015), learning that the substrate U7 residue is not essential to log-order enhancement of catalytic turnover even under physiological (therapeutic) conditions. The tetraloop capping stem II loop substantially modulates activity regardless of the presence of a U7 residue. These findings strongly suggest that the enzymatic capacity and structural dynamics of the hhRz are far from fully understood. The catalytic enhancements observed may bode very well for a revitalization of EhhRzs in the context of human nucleic acid therapeutics.

## MATERIAL AND METHODS

### Reagents

Synthetic DNA templates, primers, and RNA oligonucleotides were synthesized by IDT Technologies (Coralville, IA). RNA oligonucleotides were subjected to analytical reverse-phase high-performance liquid chromatography (HPLC) and purified by RNase-free HPLC. The lyophilized material supplied was solubilized in RNase-free double deionized water to achieve stock concentrations of 100 μM.

### *In vitro* transcription

*In vitro* transcription templates were generated by PCR using PfuUltra II. PCR reactions were carried out with 100 ng of single-stranded DNA template, 0.5μM upstream primer, 0.5μM downstream primer, and 50μl of PfuUltra II Hotstart 2× master mix (catalog number 600850; Agilent, Santa Clara, CA) in a total volume of 100μl according to the manufacturers’ recommendations. The following constant single-stranded DNA sequence includes the T7 promoter (underlined) and an upstream leader sequence: 5’-CCATGATTACGCCAAGCT<u>TAATACGACTCACTATAG</u>^+1^GG-3’. Templates generated from the T7 promoter incorporated two G’s following the G^+1^ for highly efficient transcription (Khvorova et al., 2003; Conrad et al., 2020). The constant region was extended by the following single-stranded sequences: WT 266, T<u>A</u>GAGCGTCTGATGAGGCC<u>GAAA</u>GGCCGAAAGGAAGTT; A7U 266, T<u>T</u>GAGCGTCTGATGAGGCC<u>GAAA</u>GGCCGAAAGGAAGTT; A7G, T<u>G</u>GAGCGTCTGATGAGGCC<u>GAAA</u>GGCCGAAAGGAAGTT; and A7U tetraloop cap variations, 5’- TTGAGCGTCTGATGAGGCC(N^4^)GGCCGAAAGGAAGTT; the “A7” residue and the stem II tetraloop sequences are underlined. N^4^ is not a randomized sequence but refers to the sense sequence of the various tetraloop sequences tested in this study (see **Fig. 3A, B**). The constant PCR sense primer had the following sequence: 5’-CCATGATTACGCCAAGCTTAATACG-3’. The PCR antisense primer, 5’-AACTTCCTTTCGGCCTTTCGGC-3’ was used to generate WT 266 and A7U 266 PCR products. The antisense primer for tetraloop variations was 5’-AACTTCCTTTCGGCC(N^4^)GGC-3’. PCR templates were purified using a QIAquick PCR purification kit (Qiagen, Hilden, Germany) and quantified by optical density at 260 nm (OD_260_) on a Nanodrop 2000 microfluidic UV-visible absorption spectrophotometer (Thermo Fisher Scientific, Waltham, MA).

For the HH16 hhRz experiments, the native HH16 substrate (17-mer) (Hertel et al., 1994, 1996; Hertel and Uhlenbeck, 1995) was modified by adding a 5’ A residue (to add 6-carboxyfluorescein [FAM] and avoid quenching by the 5’ G present in the 17-mer). The sequence of the HH16 substrate (18-mer) was 5’-AGGGAACGUCGUCGUCGC-3’. This substrate forms 8 nt of Watson-Crick base pairing on both flanks of the HH16 enzyme. The native 17-mer HH16 substrate had the same number of predicted base pairs (four) and structure of the 18-mer HH16 substrate and the same minimal free energy (see **Supp. Fig. 3**).

The 20-μl *in vitro* transcription reactions contained 500 ng of PCR product, 7.5 mM of each ribonucleotide triphosphate, and 2 μl T7 RNA polymerase and were incubated at 37°C for 3 h 45 min (MEGAshortscript; Invitrogen, Waltham, MA). Plasmid DNA was digested with 2 U of Turbo DNase (RNase-free; Invitrogen) for 15 min at 37°C. The RNA was cleaned using a Zymo RNA Clean & Concentrator-25 (catalog number R1017; Zymo Research, Irvine CA) according to the manufacturers’ recommendations and resuspended in RNase-free, DNase-free deionized water. The RNA concentration was quantified and quality controlled via the OD_260_ and OD_280_ measured on a Nanodrop 2000 spectrophotometer. hhRz RNAs were stored at −80°C until use, thawed at room temperature, and then maintained at 4°C until use.

### Fluorescence-based real-time ribozyme cleavage assay

Target mRNA was designed as a synthetic 15-mer human *RHO* mRNA substrate or a 14-mer *RHO* substrate containing a fluorescent dye (FAM) at the 5’ end and a quenching dye (black hole quencher 1 [BHQ-1]) at the 3’ end. The FAM is attached to the 5’ nucleotide phosphate through a 6-carbon linker (IDT). FAM has an excitation maximum of 495 nm and emission maximum 520 nm. BHQ-1 has an absorbance maximum at 534 nm; the extinction coefficient at absorbance maximum is 34,000 M^−1^ cm^−1^. All synthetic substrates were prepared by IDT, RNase-free HPLC purification, and quality controlled by the manufacturer. The binding, cleavage, and dissociation of products separates the upstream FAM-tagged fragment from the downstream BHQ1-tagged fragment to enable detection of FAM fluorescence, which is measured in real time with a Smart Cycler II thermal cycler (Cepheid Inc., Sunnyvale, CA) using appropriate excitation (LED) and emission filtering for FAM. The thermal cycler was programmed to sample the reaction at constant temperature (37°C). For short reaction cycles (300 s), the thermocycler sampled the FAM fluorescence every 12 s (6-s dwell time and 6 s of read time with the measurement made at the end of each read time cycle); for longer reaction time courses, sampling occurred every 120 s. Steady-state reactions (substrate excess) were linear in quality, and the rate of cleavage was estimated for each reaction in Excel, within the first 25 optical samples (within 300 s), based on the point of rise of FAM fluorescence (intercept) and the approach to saturation of the photodiode in the particular reaction wells used in each experiment. This fitting was conducted by eye, extending from the initial point(s) of the reaction along the linear aspect of the fluorescence increase. Steady-state (turnover) assays were conducted at a 1:10 molar ratio of hhRz to substrate and were initiated by adding the hhRz to a preformed solution containing buffer, substrate RNA, and Mg^2+^, with all components prechilled on ice. Pre-steady-state and single-turnover reactions were conducted at a 10:1 molar ratio of hhRz to substrate in 10 mM Tris-HCl buffer (pH 7.5) with pre-annealing (95°C for 2 min, 65°C for 2 min, and 27°C until reaction was started); the reaction was catalyzed by adding Mg^2+^ from a concentrated stock with rapid manual mixing and pulse centrifugation in the Cepheid cuvette. Pre-steady-state (single-turnover kinetic) reactions were fit by nonlinear least-squares minimization (Marquardt analysis).

### Gel-based cleavage assay

An *in vitro* cleavage assay was initiated by adding 100 nM EhhRz to a 25-μl reaction mixture containing 1 μM of substrate, 10 mM Tris-HCl (pH 7.5), and 0.5 mM MgCl_2_. The reaction was incubated at 37°C for 5 min and then terminated by adding 25 μl of Gel Loading Buffer II (Invitrogen) and incubating at 95°C for 5 min. Samples were run on a denaturing 12% polyacrylamide 8 M urea gel at 50 mV for approximately 3 h. The gel was removed from the box and placed in 1× Tris-borate-EDTA buffer. The FAM dye moiety at the 5’ end of either the substrate (15- or 14-mer) or the upstream cleavage product (8-mer) was visualized in the gel by using a ChemiDoc MP system (Bio-Rad, Hercules, CA) with a fluorescein filter. The gel was then poststained with SYBR Gold (Invitrogen) for 10 min to assess all RNAs in the reaction. The SYBR Gold filter on the ChemiDoc was used to visualize the hhRz, substrate, and cleavage products.

### RNA folding

Two-dimensional RNA folding was predicted with RNAStructure (v6.2) (Reuter and Mathews, 2010) using a maximum energy difference of 10%, 20 as the maximum number of structures, and a window size of 2. All suboptimal structures were generated in RNAStructure with a maximum percent energy difference of 50% and a maximum absolute energy difference of 10%. RNA Composer (Popenda et al., 2012; Antczak et al., 2016) was used to predict the three-dimensional RNA structure.

### Data analysis

Statistical analysis and linear and nonlinear functional fitting were conducted in Origin software (OriginLab Corp., Northampton, MA). The criterion for significance was a *p* value of <0.05.

## RESULTS

### High-performance hhRz kinetics assay

The 266 CUC↓ hhRz targets a region in human *RHO* mRNA accessible at both the secondary and tertiary structural levels (Abdelmaksoud et al., 2009; Yau et al., 2016; Yau et al., 2019). Here, we developed a novel moderate-throughput hhRz fluorescence cleavage assay that does not require gels (**Fig. 1A**). Fluorescence from short RNA substrates with a 5’ FAM fluorophore is partially quenched by a 3’ BHQ1 quencher because of the physical proximity in the intact substrate. Biophysically, this occurs because of Förster interactions between the fluorophore and the quencher (1/r^6^ dependence, where r is the distance between 10 and 100 Å) that result in fluorescence resonance energy transfer. Site-specific cleavage of the annealed substrate by the hhRz, followed by product release and diffusion (distancing), leads to an increase in fluorescence detectable in real time on a standard real-time PCR machine with optical detectors (e.g., silicon photodiodes) (**Fig. 1B**). The amount of FAM fluorescence signal is linear with respect to the substrate concentration for 15-mer and 14-mer substrates and for the 8-mer fluorescent 5’ product (**Supp. Fig. 1**); each cleaved and released substrate molecule is expected to contribute linearly to the fluorescence emission signal, and 8-mer product fluorescence is estimated at 3.11 FAM U nM^−1^. Initial rates were estimated by fitting a line by eye to the initial data points in the time-dependent fluorescence emission data, which corresponded to rates estimated by linear least-squares fitting, thereby validating the data reduction process (**Fig. 1B**). The 15-mer substrate alone showed steady background fluorescence over time. EhhRzs harboring the wild-type stem-loop II tetraloop [WT(GAAA)], and those harboring the altered U7 residue and either wild-type or altered tetraloop [A7U(GAAA) or A7U(AGUA), respectively] had progressively faster rates of turnover originating from the substrate-alone baseline. The positive slopes are the result of catalytic turnover, given that catalytic core mutations (G5C, G8C, G12C, and aggregated or independent mutations [not all shown]) known to prevent catalysis (Ruffner et al., 1990; Hertel et al., 1992) obviate optical turnover measure [here shown for the A7U(GAAA) construct]. Also, substrates with a non-cleavable CUG triplet showed no evidence of turnover. HH16, a well-studied mhhRz with a single-turnover rate of up to 1 min^−1^ at supraphysiological Mg^2+^ levels (≥10 mM) and lower temperature (∼25°C) demonstrated a slow steady-state cleavage rate in this assay with a labeled 18-mer substrate over a 5-min reaction under conditions of 0.5 mM Mg^2+^ and 37°C; the lower level of baseline fluorescence with the HH16 reaction is likely an index of quenching as a result of substrate structure (see below). Plotting of HH16 product fluorescence versus the concentration gave an estimated slope of 0.98 FAM U nM^−1^ (**Supp. Fig. 2**). Estimated turnover rate, corrected for product fluorescence and the optical effects of substrate binding, is approximately 9 nM min^-1^.

**Figure 1.**
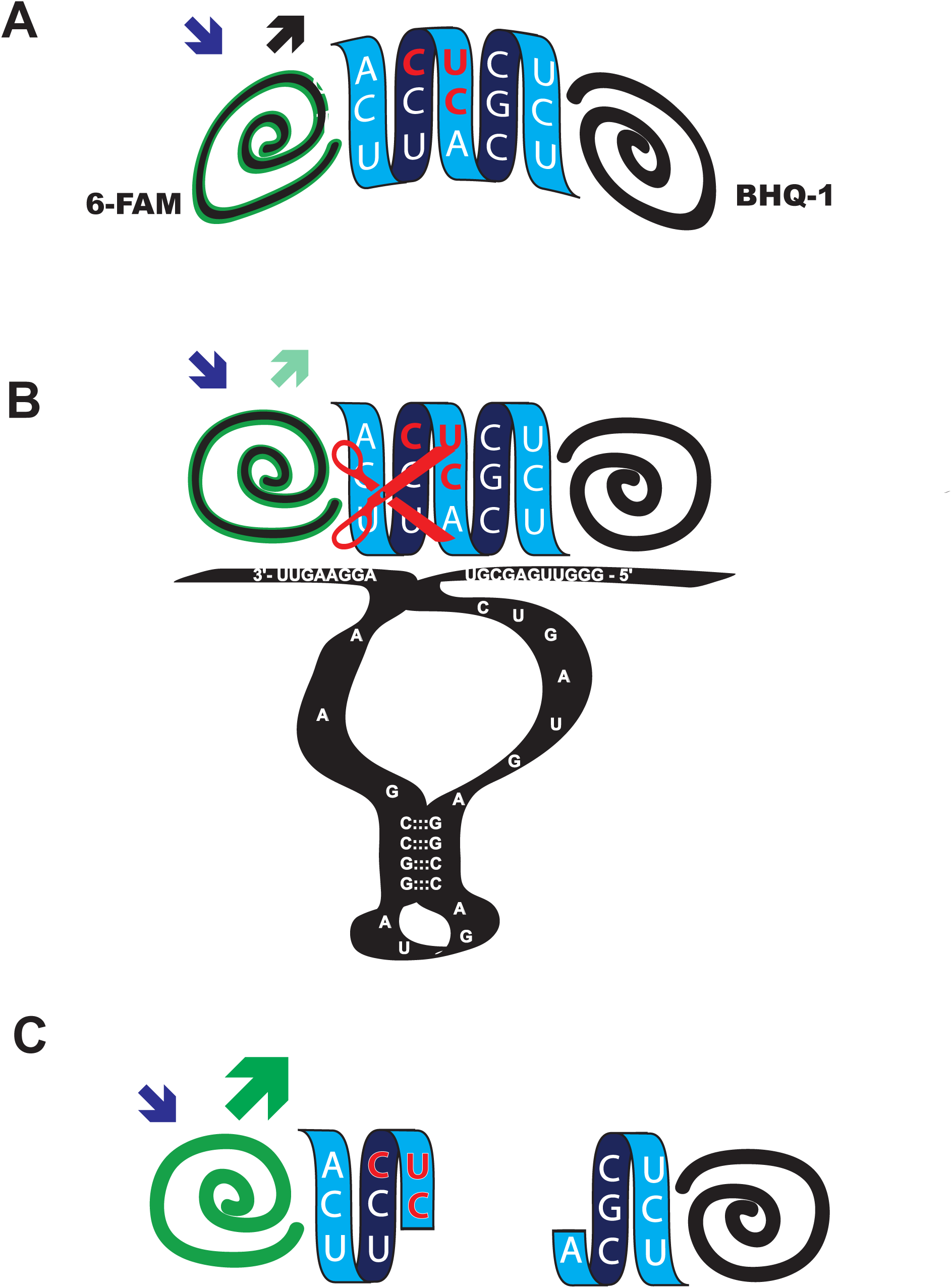
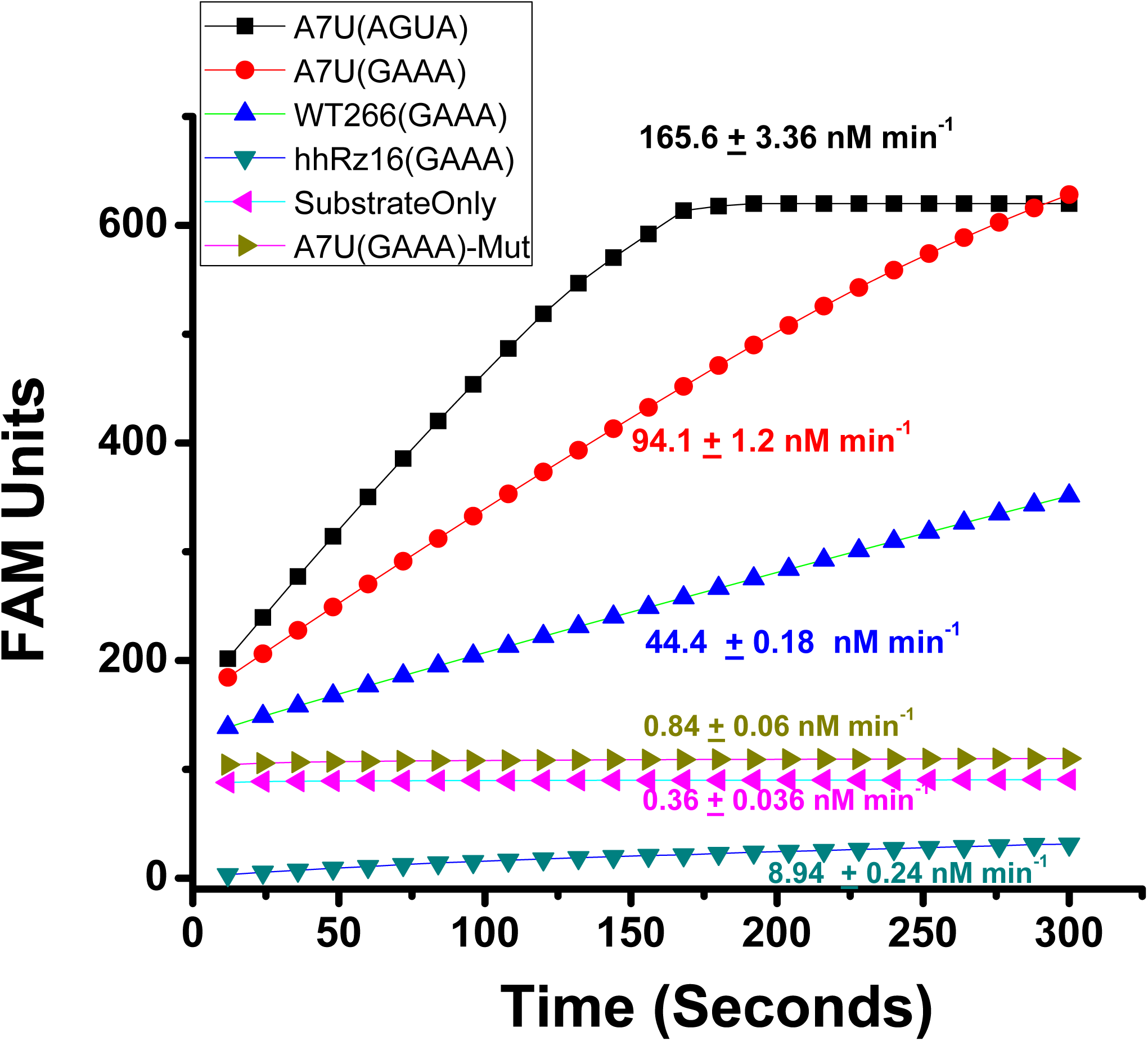
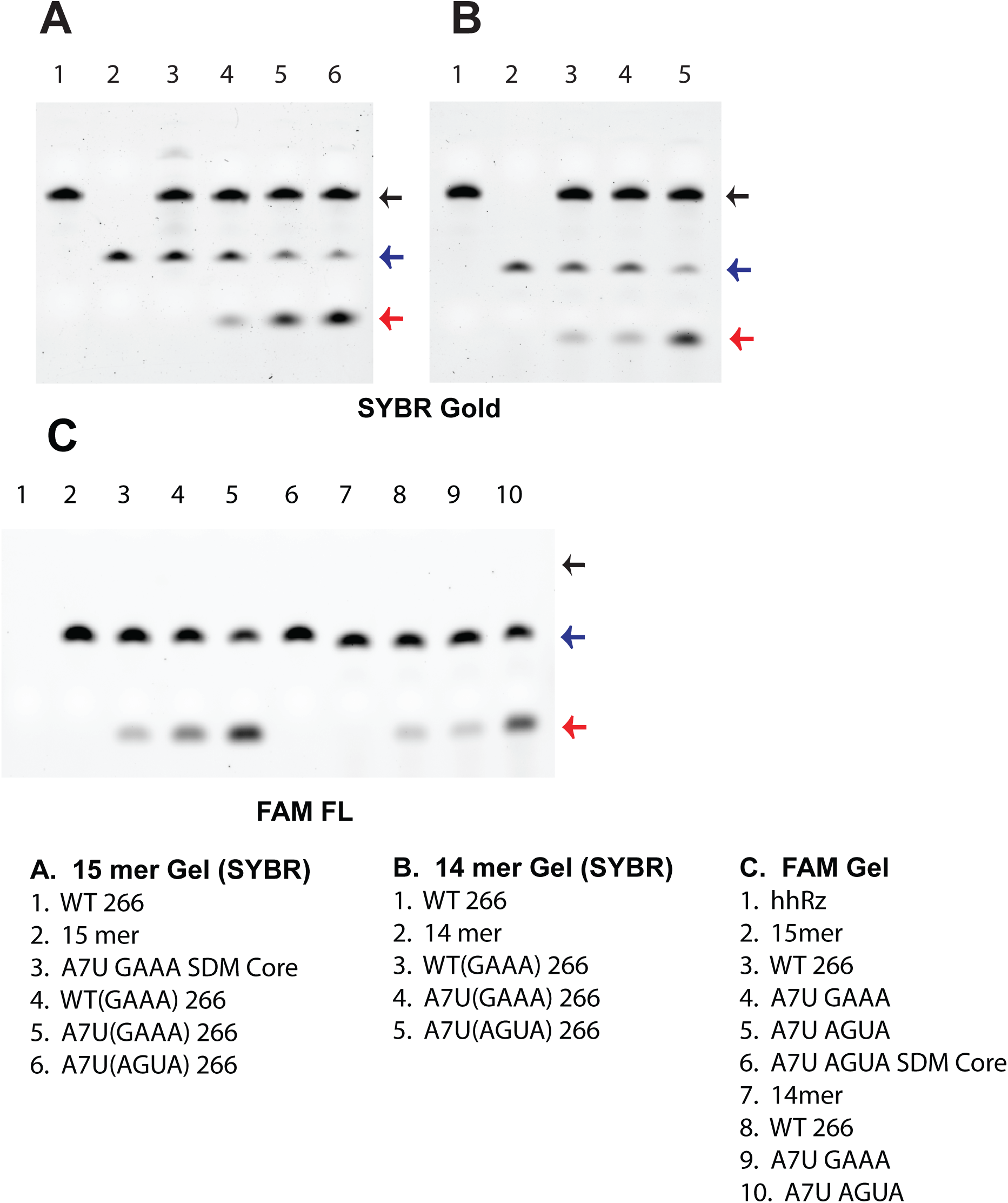
Quantitative high-throughput fluorescence assay for hhRz kinetics. (**A, a**) The short RNA substrate has a fluorophore (here, FAM) at the 5’ end and a quencher (here, black hole quencher 1 [BHQ-1]) at the 3’ end. The short substrate is drawn as a random coil (slight arc) without significant secondary structure. The quencher and the fluorophore are within Förster distance (1/r^6^ dependence, 10–100 Å with R_0_ [50%] distance particular to each pair) such that the fluorescence energy emitted from FAM is absorbed (quenched) by BHQ-1. (**A, b**) Upon annealing of the hhRz, the random coil of the substrate is disrupted with further separation from the quencher, thus increasing fluorescence emission. (**A, c**) Upon cleavage of the substrate by the hhRz at the NUH↓ site (here CUC↓), release of products, and diffusion, the fluorophore becomes well separated from the quencher, and emission is significantly enhanced (shown in schematic as increased emission arrow). This generates a real-time quantitative assay of enzymatic turnover (binding, cleavage, product release → next cycle), which can be managed in a quantitative PCR machine. (**B**) Time-dependent accumulation of FAM fluorescence in hhRz reactions. All reactions were conducted with substrate in 10-fold molar excess over the hhRz (1 μM and 0.1 μM, respectively), with 0.5 mM Mg^2+^ at 37°C. Reactions were initiated by adding the hhRz to the substrate equilibrated in buffer (10 mM Tris-HCL, pH 7.5). The initial slopes of the time-dependent FAM fluorescence changes were used in subsequent analyses. Results were scaled by the FAM units per nanomolar for the product (without BHQ1). WT(GAAA) 266 is the first EhhRz identified and showed a significant positive slope well fit by a linear function. A7U(GAAA) is the WT 266 EhhRz with the upstream “A7” residue of the antisense flank mutated to U (to prevent base pairing with substrate at U7) and showed a stronger positive slope, also well fit by a linear function. A7U(AGUA) is the A7U 266 EhhRz with the stem II tetraloop changed from GAAA to AGUA and is the fastest EhhRz that we have identified to date; it ultimately saturated the silicon photodiode detector in the qPCR machine. The substrate alone showed no definitive positive rate. Mutation of the catalytic core (G5C, G8C, G12C) (Ruffner et al., 1990) in A7U(GAAA) EhhRz also obviated the turnover activity of the enzyme. HH16 is a well-studied mhhRz (Hertel et al., 1994, 1996) and showed a slight fluorescence change during the 5-min recording cycle. Note that all the active EhhRzs showed a positive fluorescence shift evident above that for the substrate alone at time zero; we expect that this relates to the binding of the substrate by the enzyme. (**C**) Gel-based analysis corresponding to the optical assay. SYBR Gold staining for gels with the 15-mer substrate (**C, a**) and the 14-mer substrate (**C, b**) showed the definitive formation of product(s) of expected size during the 5-min reaction with substrate in excess over EhhRz (10:1 molar ratio) and 0.5 mM Mg^2+^. (**C, c**) Conversion of substrate to product is apparent in the gel imaged for FAM fluorescence. Catalytic core mutation in the EhhRzs completely suppressed cleavage, as expected. Black arrows, EhhRz; blue arrows, FAM-labeled substrate; red arrows, FAM-labeled product.

All active 266 *RHO* EhhRzs in this study drove substrate cleavage to completion. In classic gel-based assays, the EhhRzs cleave 15-mer and 14-mer substrates to yield the quantitative conversion of substrate to products within a 5-min reaction time; the products comigrate and are observable as a single band on PAGE urea gels when imaged for FAM and then poststained and imaged with SYBR Gold (RNA backbone fluorophore) (**Fig. 1C**). These PAGE results validate this moderate-throughput optical assay. The enzyme core of the EhhRzs developed here was based on the enzyme core, stem-loop II, and cap (GAAA) of the HH16 enzyme, which was extensively investigated by Uhlenbeck’s group (Hertel et al., 1994, 1996, 1997; Hertel and Uhlenbeck, 1995). We note that the 15-mer 266 *RHO* substrate has no secondary or tertiary structure in predictive algorithms (MFold and RNA Composer; **Supp. Fig. 3A–C**). By contrast, the HH16 substrate (18-mer) has strong secondary structure (single structure predicted with Δ*G* = -1.3 kCal mol^−1^; **Supp. Fig. 3D–F**) and does not possess a U7 residue 3’ of the GUC↓ cleavage motif.

### U7 substrate residue of the “uncompromised” mhhRz is not essential

Evidence from natural occurring extended hhRzs acting *in cis* and engineered hhRzs designed for *in trans* function indicates that the stem II loop of hhRz interacts with the upstream region of hhRz while bound to its downstream substrate flank (Khvorova et al., 2003; De La Peña et al., 2003; Peneda et al., 2004). We sought to investigate the expected interaction between the upstream-bound antisense flank and the loop capping stem II of the hhRz core. In *RHO* mRNA there is a U residue at the seventh nucleotide position (underlined) downstream of the substrate cleavage site after position 266 (5’-AAUUCCUC↓ACGCUC<u>U</u>-3’). O’Rourke et al. (2015) found an *unpaired* U7 residue forming a Hoogsteen base pair interaction with the fourth nucleotide of the GUG<u>A</u> stem II cap in their hhRz:substrate complex. They proposed that this interaction is *essential* and *sufficient* to generate high hhRz enzyme activity, specifically in their assays under enzyme excess (pre-annealed) and high Mg^2+^ (10 mM) conditions. In our hands, under multi-turnover conditions ([S]/[E] = 10:1 [1 μM substrate to 0.1 μM EhhRz 266]) and at physiological intracellular concentrations of free Mg^2+^ (0.5 mM) and temperature (37°C), WT EhhRz 266 showed strong levels of activity (mean ± SD): 46.48 ± 5.42 (SD) min^−1^, *n* = 36), greater than the rates expected for an mhhRz (**Fig. 2A**). We hypothesized that the U7 residue of the substrate, being paired with a Watson-Crick complementary “A7” on the upstream antisense flank of the mhhRz, is partially constrained from forming a Hoogsteen face base pairing with the A_4_ residue of the GAAA stem II loop. An A7U mutation in the upstream antisense flank of the EhhRz eliminates this constraint on the substrate U7 residue by obviating the Watson-Crick interaction. The turnover rate of A7U EhhRz 266 was significantly enhanced relative to that of the WT enzyme (96.84 ± 18.21 min^−1^, *n* = 52). By contrast, when the Watson-Crick base at the same position is maintained with an A7G mutation in EhhRz (U:A has free energy similar to that of a U:G base pair), the rate was significantly decreased relative to those of both the WT and A7U EhhRzs (35.55 ± 3.96 min^−1^, *n* = 23). Statistical comparison by one-way ANOVA across all samples showed significant differences (*p* = 3.86 E-41). Bonferroni testing showed the turnover rates of A7U (*p* = 1.59E−33) and A7G (*p* = 0.006) EhhRzs were significantly different from that of the WT and from each other (*p* = 2.16E−35).

**Figure 2.**
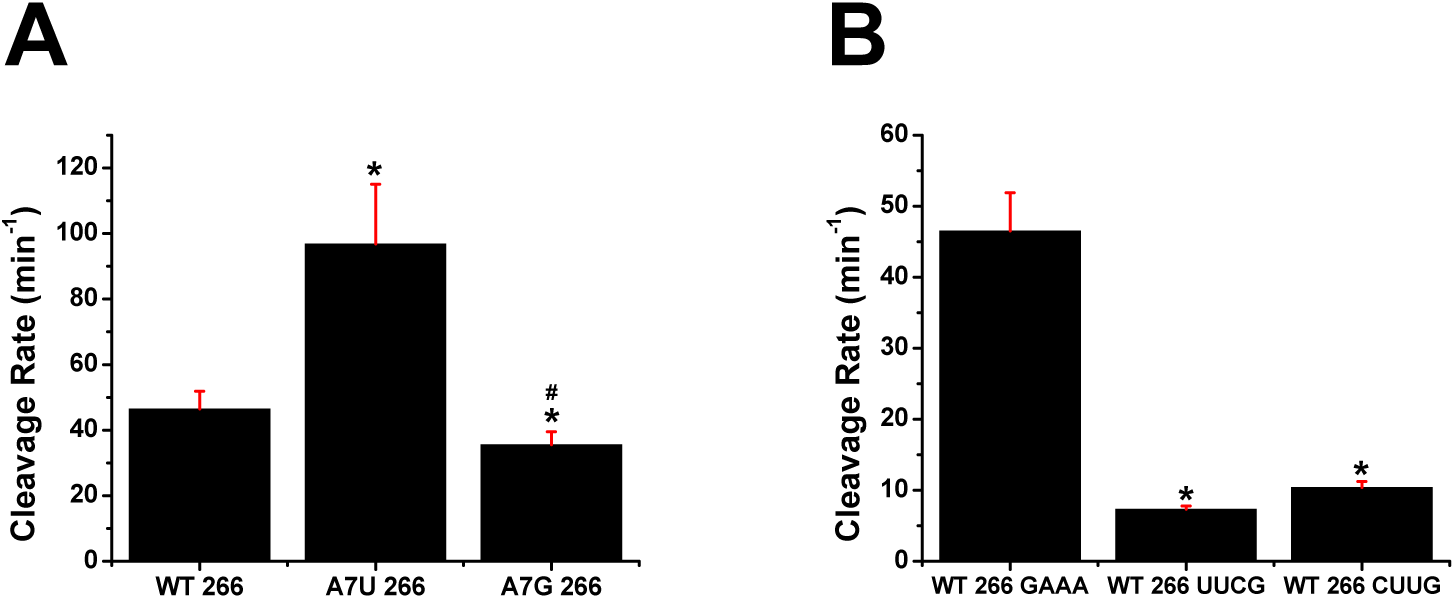
Comparison of kinetic impacts of hhRz upstream A7 replacements and stem II tetraloop replacements. Means and standard deviations are shown in the graphs. (**A**) Upstream A7 replacements. The substrate U7 residue in our WT(GAAA) EhhRz is in a Watson-Crick base pair with “A7” in the upstream antisense flank of the EhhRz. We use the nomenclature that the A7 residue of the hhRz is the nucleotide in the *upstream* antisense flank that pairs with the U7 in the substrate RNA. U7 is expected to interact with the fourth nucleotide (underlined) of the GAA<u>A</u> tetraloop triplet of the hhRz (O’Rourke et al., 2015) via Hoogsteen face interactions. All rates were scaled by product fluorescence per nanomolar (3.11 FAM units/nM). We mutated the A7 residue in the hhRz to U (A7U) to obviate formation of the Watson-Crick base pair with the substrate to relieve the constraint that could influence interactions with the tetraloop. We also mutated the hhRz A7 residue to G (A7G), which can still interact with substrate U7 by Watson-Crick pairing (G:U). Sample numbers were as follows: WT 266, *n* = 36; A7U 266, *n* = 52; A7G 266, *n* = 23. Kolmogorov-Smirnov tests for normality indicated that the samples were drawn from a normally distributed populations. Homogeneity of variance test (Levene’s absolute) showed significant differences in population variance (*p* = 1.69E-11). By parametric one-way ANOVA the sample means were found to be different (*p* = 3.86E-41). Post-hoc testing by both parametric and nonparametric analyses indicated that the WT differs (\**p* < 0.05) from A7U (*p* values: Bonferroni, 1.59E−33; Tukey, <0.01; Fisher, 5.31E−34; Bonferroni-Holm, 5.31E−34) and A7G (*p* values: Bonferroni, 0.0064; Tukey, 0.006; Fisher, 0.002; Bonferroni-Holm, 7.21E-36), and that the A7U and A7G variants also have different means (^#^*p* < 0.05) (*p* values: Bonferroni, 2.16E−35; Tukey, <0.01; Fisher, 7.21E−36; Bonferroni-Holm, 7.21E−36). By nonparametric Kruskal-Wallis ANOVA the sample means were found to be different (Chi-square p= 1.33E-20) and post-hoc Dunn’s Test found WT different from A7U (p=2.64E-10) and A7G (p= 0.007) and A7U different from A7G (p=2.12E-18). (**B**) Tetraloop replacements. The activity of WT(GAAA) EhhRz was compared to that of the WT(UUCG) and WT(CUUG) enzymes with stem II tetraloop variations. GAAA is from the class of GNRA tetraloops, and UUCG is from the class of UNCG tetraloops. Sample numbers were as follows: WT 266(GAAA), *n* = 36; WT 266(UUCG), *n* = 8; WT 266(CUUG), *n* = 8. Kolmogorov-Smirnov tests for normality indicated that the samples were drawn from normally distributed populations. Homogeneity of variance test (Levene’s absolute) showed significant differences in population variance (*p* = 4.58E-6). By parametric one-way ANOVA the sample means were found to be different (*p* = 2.27E-30). Post-hoc testing by both parametric and nonparametric analyses indicated that the WT 266(GAAA) differs (\**p* < 0.05) from WT 266 (UUCG) (*p* values: Bonferroni, 2.08E−26; Tukey, <0.01; Fisher, 5.31E−34; Bonferroni-Holm, 6.93E−27) and from WT 266(CUUG) (*p* values: Bonferroni, 7.55E-25; Tukey, <0.01; Fisher, 2.52E-25; Bonferroni-Holm, 2.52E-25); the WT 266(UUCG) and WT 266(CUUG) means were not different (*p* values: Bonferroni, 0.572; Tukey, 0.38; Fisher, 0.19; Bonferroni-Holm, 0.19). By nonparametric Kruskal-Wallis ANOVA the sample means were found to be different (Chi-square p= 4.71E-8) and post-hoc Dunn’s Test found WT 266(GAAA) different from WT 266(UUCG) (p= 1.22E-6) and WT 266(CUUG) (p= 6.09E-4) but WT(UUCG) not different from WT(CUUG) (p= 0.87).

With the substrate U7 paired to the complementary “A7” in the WT EhhRz (46.48 ± 5.42 min^−1^, n=36), we mutated the GAAA tetraloop capping stem II to a highly stable UUCG tetraloop, which dramatically reduced turnover activity (7.32 ± 0.46 min^−1^, Bonferroni *p* = 2.08E−26 versus WT) (**Fig. 2B**). An alternative CUUG had a similar impact (10.38 ± 0.86 min^−1^, Bonferroni *p* = 7.55 E−25 versus WT); the turnover rates of EhhRzs with UUCG and CUUG tetraloops were not significantly different (*p* = 0.57). These outcomes confirm that stem II loop interacts with the upstream region of the hhRz in the EhhRz format (O’Rourke et al., 2015; Khvorova et al. 2003). The physical chemical properties (i.e., energy and structure) of the Watson-Crick base pair of the hhRz with the substrate U7 residue influence the interaction with stem-loop II, and this substantially impacts the turnover rate. The interaction is optimized when the U7 substrate residue is not base paired to the EhhRz.

To further investigate the stem II loop interaction, we varied the stem II cap sequence from the wild type (“WT”) 4-nt GAAA tetraloop to a variety of alternatives in the A7U EhhRz (23 total; **Fig. 3A**). The standard 15-mer substrate was used under multi-turnover conditions ([S]>>[E]; 1:0.1 μM) at physiological [Mg^2+^] (0.5 mM) and temperature (37°C) in 10 mM Tris-HCl (pH 7.5). The WT loop is a standard GNRA tetraloop (where N is any nucleotide and R is a purine), and five of our constructs were of this variety. The A7U EhhRz with the GAAA tetraloop had a turnover rate of approximately 97 min^-1^. The best tetraloops all had turnover rates greater than 150 min^-1^ and were AGUA (169.32 ± 19.86 min^-1^), AUUA (164.92 ± 31.71 min^-1^), AAUA (159.52 ± 27.80 min^-1^), and AAAA (152.40 ± 13.71 min^-1^). The AUUA, AAUA, and AAAA constructs had means that were not significantly different from AGUA (**Fig. 3A**); the AGUA construct was used for more detailed studies. Although this screen was not nearly an exhaustive search (9% of 256 [4^4^] possibilities), the outcomes demonstrate a wide range of rates with both strong enhancement and suppression of an already enhanced turnover rate observed with the WT A7U EhhRz 266: the minimum with CUUG (6.63 ± 0.85 min^−1^) ≤ WT(GAAA) (96.84 ± 18.21 min^−1^) ≤ maximum with AGUA (169.32 ± 19.86 min^−1^). A one-way ANOVA refuted the null hypothesis of equivalence across the ensemble of constructs (*F* = 70.79, *p* = 1.225 E-114). Internal pairwise comparisons to the WT(GAAA) construct with parametric and nonparametric tests showed significant differences (*p* < 0.05, Bonferroni testing, asterisks) between the WT tetraloop (GAAA) and the alternatives, as evaluated by multiple statistical testing comparisons (data available upon request). These results show the extensive effect of varying the stem II capping tetraloop on EhhRz catalytic activity (13 of the 22 tetraloop variants [59%] were significantly different than the GAAA control, with some differences exceeding log-order). Statistical comparison of the EhhRz with the AGUA tetraloop found statistical significance (#, Bonferroni, p< 0.05) in 17 of 22 variants (77.3%). The set of tetraloops of EhhRzs with turnover activity not statistically different from AGUA were AAAA, AUUA, AAUA, AGCA, and GAUA.

**Figure 3.**
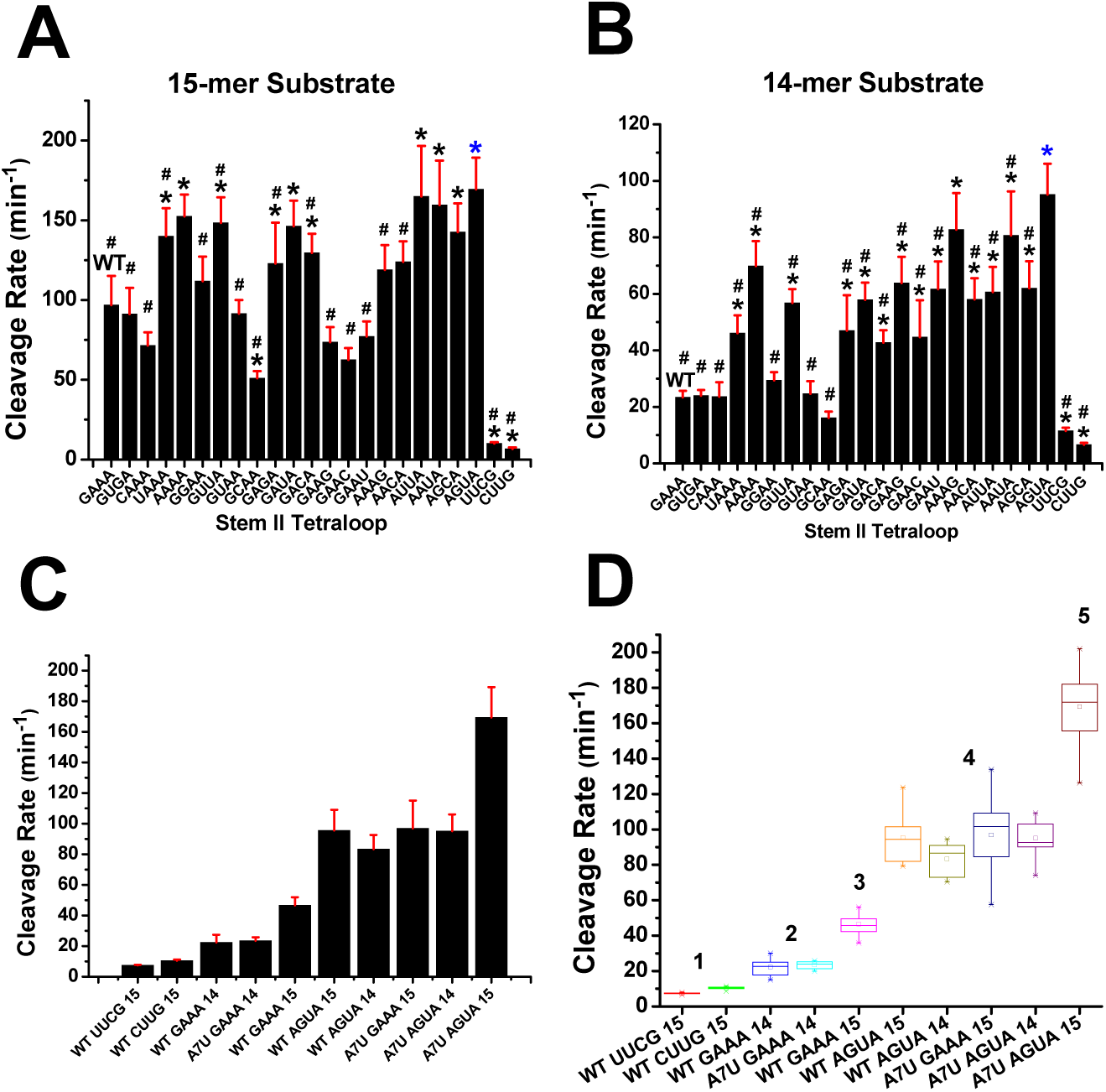
Effects of various hhRz stem-loop II tetraloop variants on catalytic activity with and without U7 in substrate. The initial rates of reaction were measured at an enzyme concentration of 100 nM and substrate concentration of 1 μM (10:1) at 0.5 mM Mg^2+^ and 37°C. All rates were scaled by product fluorescence per nanomolar (3.11 FAM units/nM). Tetraloops were varied on the 266 A7U(GAAA) EhhRz (“WT”). (**A**) 15-mer substrate RNA. Kolmogorov-Smirnov tests for normality indicated that all the samples were drawn from normally distributed populations. Homogeneity of variance test (Levene’s absolute) showed significant differences in population variance (*p* = 3.05E-13). The 266 A7U EhhRz with a WT stem II tetraloop (GAAA) showed strong activity (96.84 ± 18.21 min^-1^). When GAAA was replaced with 22 other tetraloops, the turnover activity was suppressed or enhanced relative to that with the WT tetraloop. By parametric one-way ANOVA the sample means were found to be different (*p* = 1.22E-114). Post-hoc Bonferroni tests (\**p* < 0.05) indicated means significantly different from A7U(GAAA) (“WT”). The EhhRz with greatest turnover rate relative to GAAA had an AGUA tetraloop (169.32 ± 19.86 min^-1^) (blue asterisk). Post-hoc Bonferroni tests showed constructs with turnover rate significantly different from AGUA (^#^p <0.05). Constructs with means *not* significantly different from AGUA were AAAA, GAUA, AUUA, AAUA, and AGCA. By nonparametric Kruskal-Wallis ANOVA the sample means were found to be different (Chi-square p= 3.56E-50). Post-hoc Dunn’s Test found the following constructs to have significant differences of mean vs A7U(GAAA) (“WT”): AAAA (p= 0.016), GUUA (p= 3.80E-6), AUUA (p= 7.05E-13), AAUA (p= 3.97E-9), and AGUA (p= 3.52E-16). (**B**) 14-mer substrate RNA (no U7 residue). Activity of the WT enzyme remained high (23.42 ± 2.27 min^-1^) but varied with the different stem II tetraloop compositions. Kolmogorov-Smirnov tests for normality indicated that all the samples were drawn from normally distributed populations. Homogeneity of variance test (Levene’s absolute) showed significant differences in population variance (*p* = 2.00E-15). By parametric one-way ANOVA the sample means were found to be different (*p* = 2.35E-120). Post-hoc Bonferroni tests (\**p* < 0.05) indicated means significantly different from A7U(GAAA) (“WT”). The EhhRz with greatest turnover rate relative to GAAA also had an AGUA tetraloop (95.15 ± 10.89 min^-1^) (blue asterisk). Post-hoc Bonferroni tests showed constructs with turnover rate significantly different from AGUA (^#^p <0.05). Constructs with means *not* significantly different from AGUA were only AAAG. By nonparametric Kruskal-Wallis ANOVA the sample means were found to be different (Chi-square p= 3.73E-43). Post-hoc Dunn’s Test found the following constructs to have significant differences of mean vs A7U(GAAA) (“WT”): AAAA (p= 0.003), GAUA (p= 0.026), GAAG (p= 0.032), GAAU (p= 0.006), AAAG (p= 1.54E-4), AUUA (p= 0.0095), AAUA (p= 1.00E-4), and AGUA (p= 3.16E-5). (**C**) Ordered comparison of mean rates for selected EhhRzs against 14-mer and 15-mer substrates. (**D**) Boxcar comparison of selected EhhRzs against 14-mer and 15-mer substrates shown in (C). The box outlines embrace the 75^th^ and 25th percentiles, and enclose the means (ÿ), and median (−—); the maximum and minimum values (×) and the 1^st^ and 99^th^ percentiles (−) are shown. The numbers (1-5) refer to apparent quantized presentation of outcomes.

To determine the extent to which the U7 residue impacts rate enhancement, we shortened the substrate to 14 nt by deleting the U7 residue identified by O’Rourke et al. (2015) as critical to the interactions. This change would also increase the product leaving rate of one antisense flank, which is expected to enhance the turnover rate if product release is rate limiting at 37°C in the overall mechanism (Stage-Zimmermann and Uhlenbeck, 1998). Using the same set of cap tetraloop variants, we were surprised to find a somewhat similar activity landscape (**Fig. 3B**). The four best tetraloops all had turnover rates greater than 65 min^-1^ and were AGUA (95.15 ± 10.89 min^-1^), AAAG (82.78 ± 12.90 min^-1^), AAUA (80.70 ± 15.61 min^-1^), and AAAA (69.85 ± 8.84 min^-1^), which were all significantly greater than GAAA. Rates with the 14-mer were uniformly lower than with the 15-mer substrate under identical reaction conditions: minimum with CUUG (6.59 ± 0.71 min^−1^) ≤ GAAA (23.42 ± 2.27 min^−1^) ≤ AGUA (95.15 ± 10.89 min^−1^). A one-way ANOVA refuted the null hypothesis of equivalence across the ensemble of constructs (*F* = 98.04, *p* = 2.35E-120). Internal pairwise comparisons to the WT(GAAA) construct showed significant differences (*p* < 0.05, Bonferroni testing) for many constructs (asterisks) (statistical data available upon request for full parametric/nonparametric testing). The tetraloop of the only EhhRz with turnover activity not statistically different from that with AGUA was AAAG (p = 0.56). These results show the extensive effect of varying the stem II capping tetraloop on EhhRz catalytic activity (17 of the 22 tetraloop variants [77%] were significantly different than the GAAA control).

The best tetraloop performers were AGUA and AAAG for the 14-mer and AGUA and AUUA for the 15-mer substrates, respectively. The CUUG tetraloop yielded the lowest turnover rate for both 14-mer and 15-mer substrates. With the same capping tetraloop, predicted improvements in the leaving rate for the 14-mer substrate were expected to enhance the overall turnover rate rather than decrease it. Calculations of the equilibrium constants and rates for the hhRz reaction scheme, by way of an established nearest-neighbor model approach (Stage-Zimmermann and Uhlenbeck, 1998), confirmed this (**Supp. Table 1**). For a given substrate, the enhancement or suppression of turnover rate must relate to structural and chemical interactions that drive the formation of the catalytically active state(s) leading to target cleavage (*k*_2_). Overall, major findings are that the composition of the tetraloop has a marked impact on kinetic turnover rate, and that the U7 residue of the substrate is *not essential* to enhanced rates, in contrast to the postulate by O’Rourke et al. (2015) that it was *necessary* and *sufficient*. In the absence of the U7 in the substrate, a U7 residue in the hhRz (i.e., A7U) yields greatly enhanced rates that vary with different stem II tetraloops.

We compared the kinetic outcomes between the 15-mer and 14-mer substrates with the WT(GAAA), A7U(GAAA), and A7U(AGUA) EhhRzs (**Fig. 3C**); one-way ANOVA revealed that the differences between the means were significant (*F* = 283.31, *p* = 8.80 E-106). The addition of A7U to the WT EhhRz (GAAA) (non-pairing U7 substrate) enhanced the rate only for the 15-mer substrate. The AGUA tetraloop enhanced the activity relative to that of the WT construct (GAAA) when the U7 of the substrate was either base paired (15-mer) or absent (14-mer) and further enhanced the A7U EhhRz variation but only with the 15-mer substrate. Curiously, the activity level of WT EhhRz (GAAA) with the 14-mer was approximately one-half that of WT EhhRz (GAAA) with the 15-mer substrate, but the A7U shift did not enhance activity of the EhhRz (GAAA) with the 14-mer substrate; the A7U shift did enhance with the AGUA tetraloop acting on the 14-mer substrate. Four constructs (WT AGUA [15-mer], WT AGUA [14-mer], A7U GAAA [15-mer], and A7U AGUA [14-mer]) generated similar levels of activity [overall ANOVA p= 0.03132 with the only significant difference in the internal comparisons was WT AGUA [14-mer] *vs.* A7U(GAAA) [15-mer] (p = 0.021)]. The A7U GAAA [15-mer]) was approximately double the activity level of the WT GAAA (15-mer) reaction. The maximum turnover level was achieved with the A7U(AGUA) EhhRz and the 15-mer substrate, which was approximately double the level for the four others that appear to aggregate in activity. This finding suggests that the activity enhancements may be quantized over at least five levels; the UUCG/CUUG constructs have approximately one-half of the activity level of other constructs at lower levels (WT(GAAA) [14-mer] and A7U(GAAA) [14-mer]) (**Fig. 3D**). Without the A7U EhhRz variation, the AGUA-containing constructs are agnostic to the length of the substrates. The A7U variation at the 5’ end of EhhRz sensitizes the turnover rate to the length of the substrate. A7U generates an unpaired (unconstrained) U residue in the 5’ end of the EhhRz, leaving the U7 at the 3’ end of the substrate unpaired. The 5’ end of the EhhRz and the 3’ end of the substrate might act in relatively independent manners in the local structural space (e.g., interacting with preferential stem II loops) to influence catalytic performance; however, the molecular/atomic interactions, topology, and dynamics that drive these activity changes are beyond the scope of this initial study.

### Kinetics analysis of 266 EhhRzs

We conducted a detailed analysis of the kinetic properties of the WT(GAAA), A7U(GAAA), and A7U(AGUA) 266 *RHO* EhhRzs. Using consistent amounts of 15-mer substrate (1 μM), increasing the amount of enzyme (0–500 nM) lead to a proportional increase in the rate of the reaction measured optically under the conditions of physiological 0.5 mM Mg^2+^ and 37°C (**Fig. 4**). Linear fits of the turnover rate to enzyme concentration gave statistically reliable functions, consistent with Michaelis-Menten enzyme function, with *R*^2^ values of 0.99963 (ANOVA *p* = 1.11E−16) for WT(GAAA) EhhRz 266 (**Fig. 4A**), 0.9977 (ANOVA *p* = 3.04E-5) for A7U(GAAA) (**Fig. 4D**), and 0.99328 (ANOVA *p* = 1.08E-5) for A7U(AGUA) (**Fig. 4G**). Similarly, when the *y* intercept was fitted according to enzyme concentration, there was also a linear relationship as expected (data not shown). We subsequently conducted a formal Michaelis-Menten analysis for the three 266 hhRzs by increasing the substrate concentration but fixing the EhhRz concentration at 50 nM (linear range) under conditions with 0.5 mM Mg^2+^ at 37°C. The WT(GAAA) enzyme data fit well with the Michaelis-Menten hyperbolic function (ANOVA *p* = 1.80E−11) (**Fig. 4B**), and a fit of the Eadie-Hofstee linear transform confirmed similar parameters (ANOVA *p* = 7.97E−6) (**Fig. 4C**). The A7U(GAAA) and A7U(AGUA) kinetics were also well fit by the hyperbolic nonlinear function (ANOVA *p* = 2.41E−12 and 4.61E−12; **Fig. 4E and H**, respectively), with similar linear transform results (ANOVA *p* = 3.47E−6 and 0.00181; **Fig. 4F** and **I**, respectively). The kinetics results of these measures for the three 266 enzymes are shown in **Table 1**. Enzyme efficiency (*V*_max_/*K_m_*) or turnover is commonly used to compare different catalytic entities.

**Figure 4.**
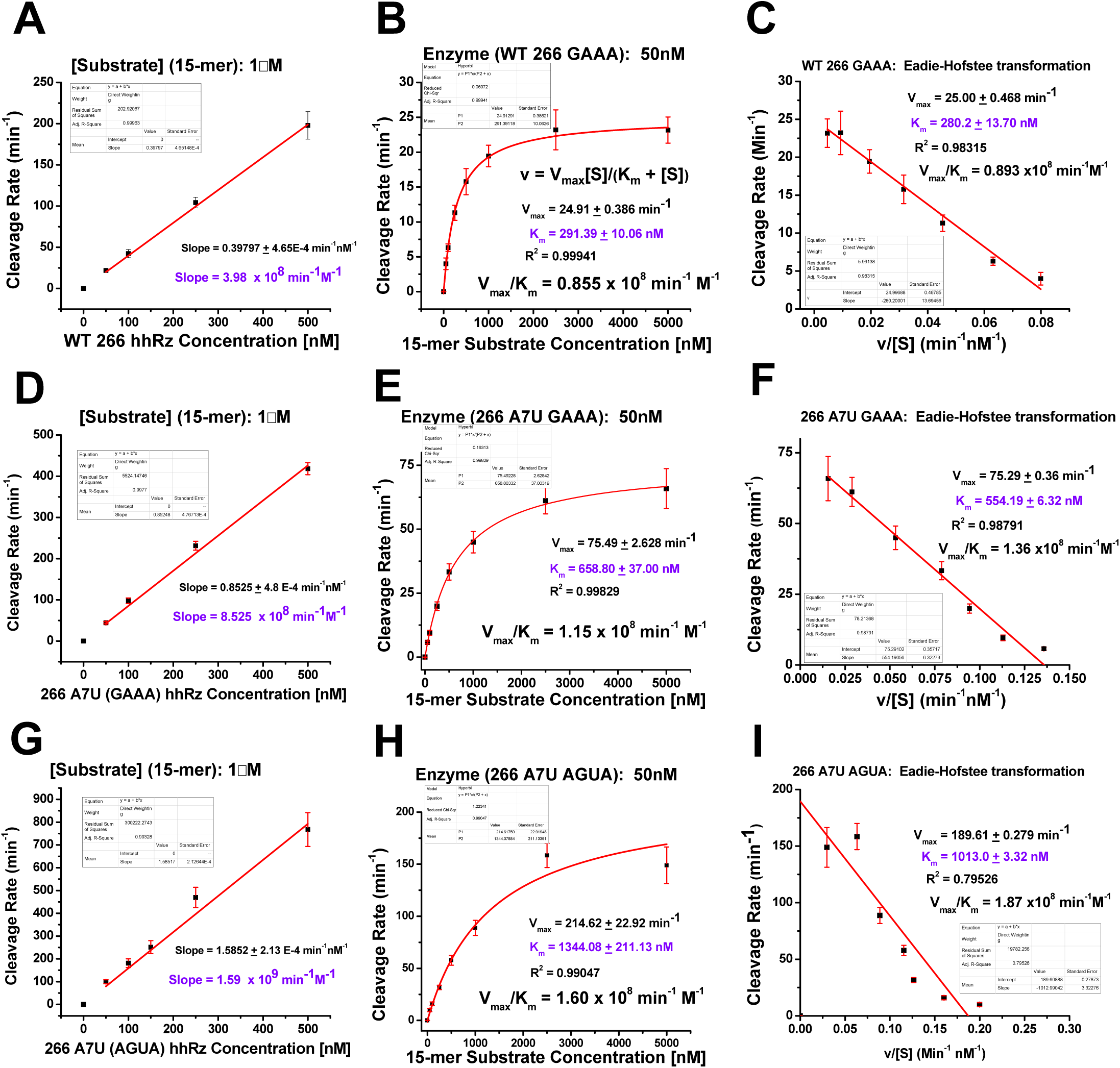
Michaelis-Menten analysis of EhhRzs against the 15-mer substrate. The turnover rates for 1μM substrate were evaluated for various concentrations of the three EhhRzs: WT(GAAA) 266 EhhRz (**A**), A7U(GAAA) 266 EhhRz (**D**), and A7U(AGUA) 266 EhhRz (**G**). All rates were scaled by product fluorescence per nanomolar (3.11 FAM Units/nM). Substrate turnover rates increased linearly over this range of enzyme concentrations for all three constructs, and the estimated turnover rates were all greater than 1 x 10^8^ min^−1^ with a rank order of A7U(AGUA) > A7U(GAAA) > WT(GAAA). Classic Michaelis-Menten analyses were conducted measuring initial rates with various substrate concentrations and the enzyme concentration fixed at 50 nM: WT(GAAA) 266 EhhRz (**B**), A7U(GAAA) 266 EhhRz (**E**), and A7U(AGUA) 266 EhhRz (**H**). All turnover titrations are well fit by the Michaelis-Menten hyperbolic function using nonlinear curve fitting. *V*_max_ and *K_m_* were parameters extracted from the fittings (**Table 1**). Rankings of *V*_max_ and *K_m_* values and enzymatic efficiency (*V*_max_/*K_m_*) (turnover) matched that for enzyme excess experiments: A7U(AGUA) > A7U(GAAA) > WT(GAAA). Eadie-Hofstee linear transforms and fitting of the Michaelis-Menten data corroborate the findings of the nonlinear curve fitting for WT(GAAA) 266 EhhRz (**C**), A7U(GAAA) 266 EhhRz (**F**), and A7U(AGUA) 266 EhhRz (**I**).

**TABLE 1.**
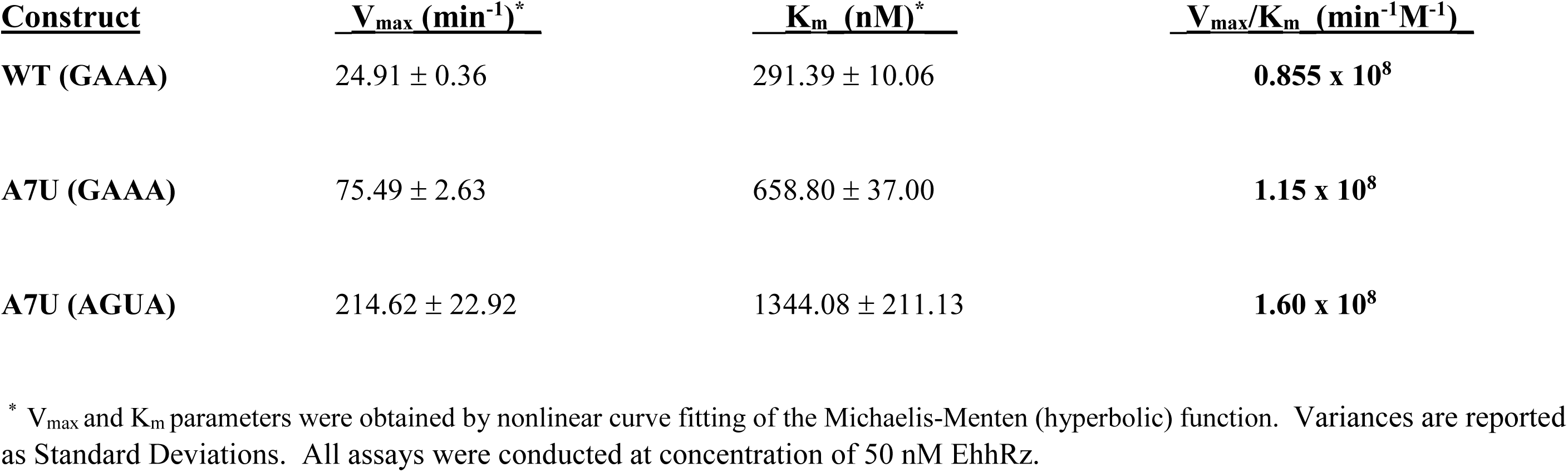

| **<u>Construct</u>** | **<u>$V_{\max}$ (min<sup>-1</sup>)\*</u>** | **<u>$K_m$ (nM)\*</u>** | **<u>$V_{\max}/K_m$ (min<sup>-1</sup>M<sup>-1</sup>)</u>** |
| --- | --- | --- | --- |
| **WT (GAAA)** | 24.91 ± 0.36 | 291.39 ± 10.06 | **0.855 x 10<sup>8</sup>** |
| **A7U (GAAA)** | 75.49 ± 2.63 | 658.80 ± 37.00 | **1.15 x 10<sup>8</sup>** |
| **A7U (AGUA)** | 214.62 ± 22.92 | 1344.08 ± 211.13 | **1.60 x 10<sup>8</sup>** |

\* $V_{\max}$ and $K_m$ parameters were obtained by nonlinear curve fitting of the Michaelis-Menten (hyperbolic) function. Variances are reported as Standard Deviations. All assays were conducted at concentration of 50 nM EhhRz.

Notably, the enzyme efficiencies of the 266 EhhRzs are on the same scale as RNase A (1.38 × 10^8^ min^−^1 M^−1^; Thompson et al., 1995), which uses the same mechanism for RNA cleavage (Emilsson et al., 2003; Breaker et al., 2003; Cochrane and Strobel, 2008). In particular, A7U(AGUA) has a V_max_ = 214.62 min^-1^, Km = 1344 nM, and V_max_/K_m_ = 1.60 x 10^8^ M^-1^ min^-1^.

### Mg^2+^ dependence of enhanced 266 hhRzs

We investigated the sensitivity of the turnover rate under [S]>>[E] conditions (1: 0.1 μM) by varying the Mg^2+^ concentration (0–20 mM) in the reaction buffer (pH 7.5 [physiological]) for the WT(GAAA), A7U(GAAA), and A7U(AGUA) 266 EhhRzs (**Fig. 5**). Given that recent work suggests there are two Mg^2+^ binding sites in the xhhRz (Mir et al., 2015; Mir and Golden 2016) and historical work suggested multiple cooperative binding sites for the mhhRz (Menger et al., 1996, 2000), we explored two types of functionality: double Boltzmann and Hill functions. The double Boltzmann (equation **1**) represents two Mg^2+^ binding sites, each with their own independent binding equilibrium.

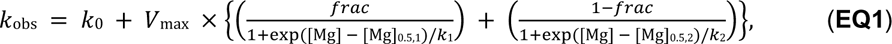

**Figure 5.**
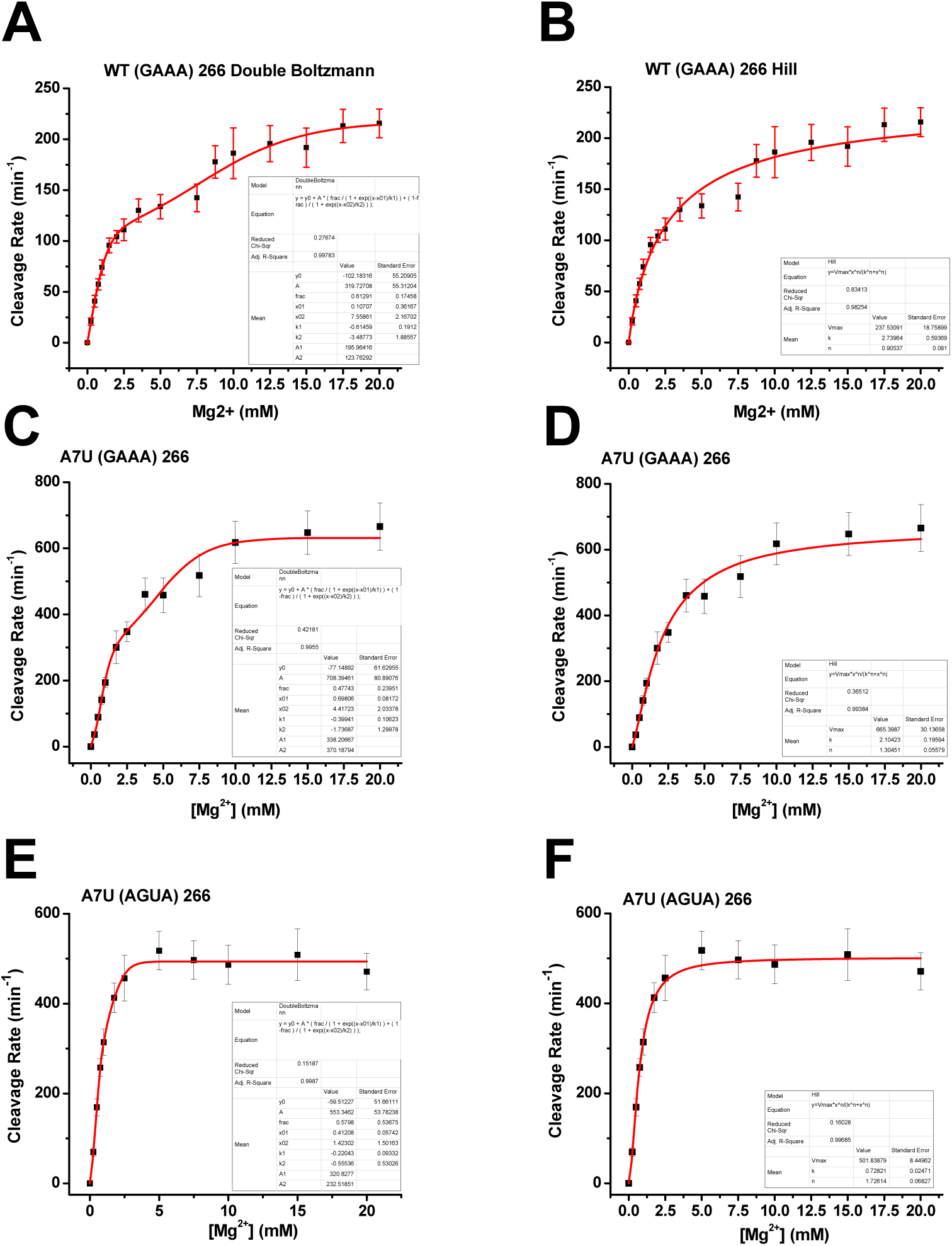
Mg^2+^ dependence of EhhRzs. Mg^2+^ concentrations were varied in reactions with 100 nM EhhRz and 1 μM 15-mer substrate at 37°C and pH 7.5 (10 mM Tris-HCl). All rates were scaled by product fluorescence per nanomolar (3.11 FAM Units/nM). The Mg^2+^ sensitivity of WT(GAAA) 266 EhhRz was well fit by a double Boltzmann function (*R*^2^ = 0.99779) (**A**) and a Hill function (*R*^2^ = 0.98254) (**B**). The Mg^2+^ sensitivity of A7U(GAAA) EhhRz was well fit by a double Boltzmann function (*R*^2^ = 0.99696) (**C**) and a Hill function (*R*^2^ = 0.99384) (**D**). The Mg^2+^ sensitivity of A7U(AGUA) EhhRz was well fit by a double Boltzmann function (*R*^2^ = 0.99522) (**E**) and a Hill function (*R*^2^ = 0.99685) (**F**). Nonlinear fitting parameters for the three EhhRzs are shown in **Table 2**.

where *k*_1_ and *k*_2_ are the sensitivity factors for the two titrations, [Mg]_0.5,1_ and [Mg]_0.5,2_ are the half points

for the independent titrations, and *frac* is the fraction of the overall fit attributed to the first (most sensitive) titration. The Hill function (Equation **2**) represents one or more cooperative Mg^2+^binding sites.

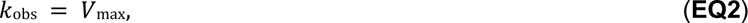

where 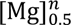 is the half point for the titration, and *n* is the cooperativity constant.

For all three EhhRzs, the turnover rate (*k*_obs_) showed a clear saturable Mg^2+^ sensitivity. Only the WT(GAAA) and A7U(GAAA) constructs showed apparent bimodal titrations. The independent site model fit the data for the WT(GAAA) (**Fig. 5A**), A7U(GAAA) (**Fig. 5C**), and A7U(AGUA) (**Fig. 5E**) EhhRzs. The Mg^2+^ isotherms were also all well fit by the Hill function for the WT(GAAA) (**Fig. 5B**), A7U(GAAA) (**Fig. 5D**), and A7U(AGUA) (**Fig. 5F**) EhhRzs. The maximum rate (*V*_max_), the *K_m_* ([Mg^2+^]_0.5_) values, and the Hill coefficients (*n*) varied with each construct (**Table 2**). For the WT(GAAA), A7U(GAAA), and A7U(AGUA) EhhRzs, the most sensitive titrations ([Mg^2+^]_0.5_ values in double Boltzmann fitting) ranged from ∼0.1 to ∼0.7 mM, and the less sensitive titrations ranged from ∼1.5 to 7.5 mM; the fractional weight of the two component titrations was approximately 50-60% for these EhhRzs. The sensitivity values for the Hill fitting for Mg^2+^ ranged from approximately 0.7 to 2.7 mM, with Hill coefficients between 0.9 (WT(GAAA) and 1.7 (A7U(AGUA)). The *V*_max_ value of the A7U(AGUA) construct is smaller than that of the A7U(GAAA) EhhRz, suggesting that the AGUA tetraloop drives highly efficient Mg^2+^-dependent formation of the most catalytically active states of the EhhRz.

**TABLE 2.** Mg^2+^ Sensitivity for EhhRzs

| **<u>Construct</u>** | **<u>V<sub>max</sub></u>** | **<u>frac</u>** | **<u>[Mg]<sub>0.5,1</sub></u>** | **<u>k<sub>1</sub></u>** | **<u>[Mg]<sub>0.5,2</sub></u>** | **<u>k<sub>2</sub></u>** | **<u>[Mg]<sub>0.5</sub></u>** | **<u>n</u>** | **<u>AdjR<sup>2</sup></u>** |
| --- | --- | --- | --- | --- | --- | --- | --- | --- | --- |
| **WT (GAAA)**<br>**Two Site** | 319.7 ±55.30 | 0.62 ±0.17 | 0.12 ±0.35 | -0.61 ± 0.19 | 7.56 ±2.17 | -3.48 ±1.89 | - | - | 0.99783 |
| **WT (GAAA)**<br>**Hill** | 237.53 ±18.76 | - | - | - | - | - | 2.74 ±0.59 | 0.91 ±0.081 | 0.98254 |
| **A7U (GAAA)**<br>**Two Site** | 708.39 ±80.9 | 0.48 ±0.24 | 0.70 ±0.08 | -0.40 ±0.11 | 4.42 ± 2.03 | -1.74 ±1.30 | - | - | 0.9955 |
| **A7U (GAAA)**<br>**Hill** | 665.40 ±30.1 | - | - | - | - | - | 2.10 ±0.20 | 1.30 ±0.06 | 0.99384 |
| **A7U (AGUA)**<br>**Two Site** | 553.3 ± 53.8 | 0.58 ± 0.54 | 0.41 ± 0.06 | -0.22 ±0.09 | 1.42 ±1.50 | -0.56 ±0.53 | - | - | 0.9987 |
| **A7U (AGUA)**<br>**Hill** | 501.84 ±8.45 | - | - | - | - | - | 0.73 ±0.02 | 1.73 ±0.07 | 0.99685 |

Nonlinear least squares fitting parameters for Mg<sup>2+</sup> sensitivity for three 266 *hRHO* EhhRzs to Equations 1 (Double Boltzmann) and 2 (Hill) in the text. **V<sub>max</sub>** is in units of **min<sup>-1</sup>**. **[Mg<sup>2+</sup>]<sub>x</sub>** parameters are in units of **mM**. **Frac**, **k<sub>1</sub>**, **k<sub>2</sub>**, and **n** are unitless. For the Double Boltzmann fit **[Mg]<sub>0.5,1</sub>** and **[Mg]<sub>0.5,2</sub>** are the Mg<sup>2+</sup> concentrations at the half maximum titration points for the separate components. For the Hill fits **[Mg]<sub>0.5</sub>** is the half-maximal titration fit. For the Double Boltzmann fitting **frac** is the Fraction of the overall **V<sub>max</sub>** that is due to the first (most sensitive) component of the Mg<sup>2+</sup> titration (1-frac is the fraction of **V<sub>max</sub>** due to the less sensitive titration). For the Double Boltzmann fits **k<sub>1</sub>** and **k<sub>2</sub>** are the sensitivity (slope) parameters for the two components of the titration. For the Hill fits **n** is the Hill exponent parameter. **AdjR<sup>2</sup>** is the adjusted R<sup>2</sup> or goodness of fit. Variances obtained from fitting are standard errors of the mean. In all cases the substrate was the 15-mer at a concentration of 1 μM and the EhhRz was at concentration of 100 nM.

### Single-turnover reaction conditions

The data presented above were collected under multi-turnover (steady state) conditions, with the substrate in excess and the reaction catalyzed by mixing the EhhRz (with Mg^2+^) with the substrate. We attempted to measure the rate of reaction at pH 7.5 with the enzyme in 10-fold molar excess over a substrate level (1 μM) that produces optical signals, with substrate and enzyme pre-annealed and with the reaction catalyzed by the addition of Mg^2+^. Under these “single-turnover” conditions, the entire population of substrate molecules is expected to be bound in Watson-Crick complex with the EhhRzs, with the cleavage reaction ready to be initiated by the Mg^2+^ cofactor (**Fig. 6**). At pH 7.5 with 0.5 mM Mg^2+^ and a 37°C reaction temperature, the initial rate of reaction for the WT(GAAA) enzyme was 273 min^−1^ (scaled) for the 15-mer substrate with a cleavable CUC↓ motif. For the A7U(GAAA) enzyme, only a few points were measurable, suggesting a rate of >680 min^−1^. The reaction was already saturated at maximum signal by the first optical measurement (12 s) for the A7U(AGUA) enzyme, suggesting a cleavage rate greater than 1000 min^-1^. As expected, there was no evidence of cleavage of the non**-**cleavable 15-mer 266 substrate (CUG) by the three EhhRzs. For all three enzymes, the large step in fluorescence before the first optical measure is an index of very rapid cleavage processes that are not captured at this kinetic bandwidth; a rapid stopped-flow kinetics analysis is necessary.

**Figure 6.**
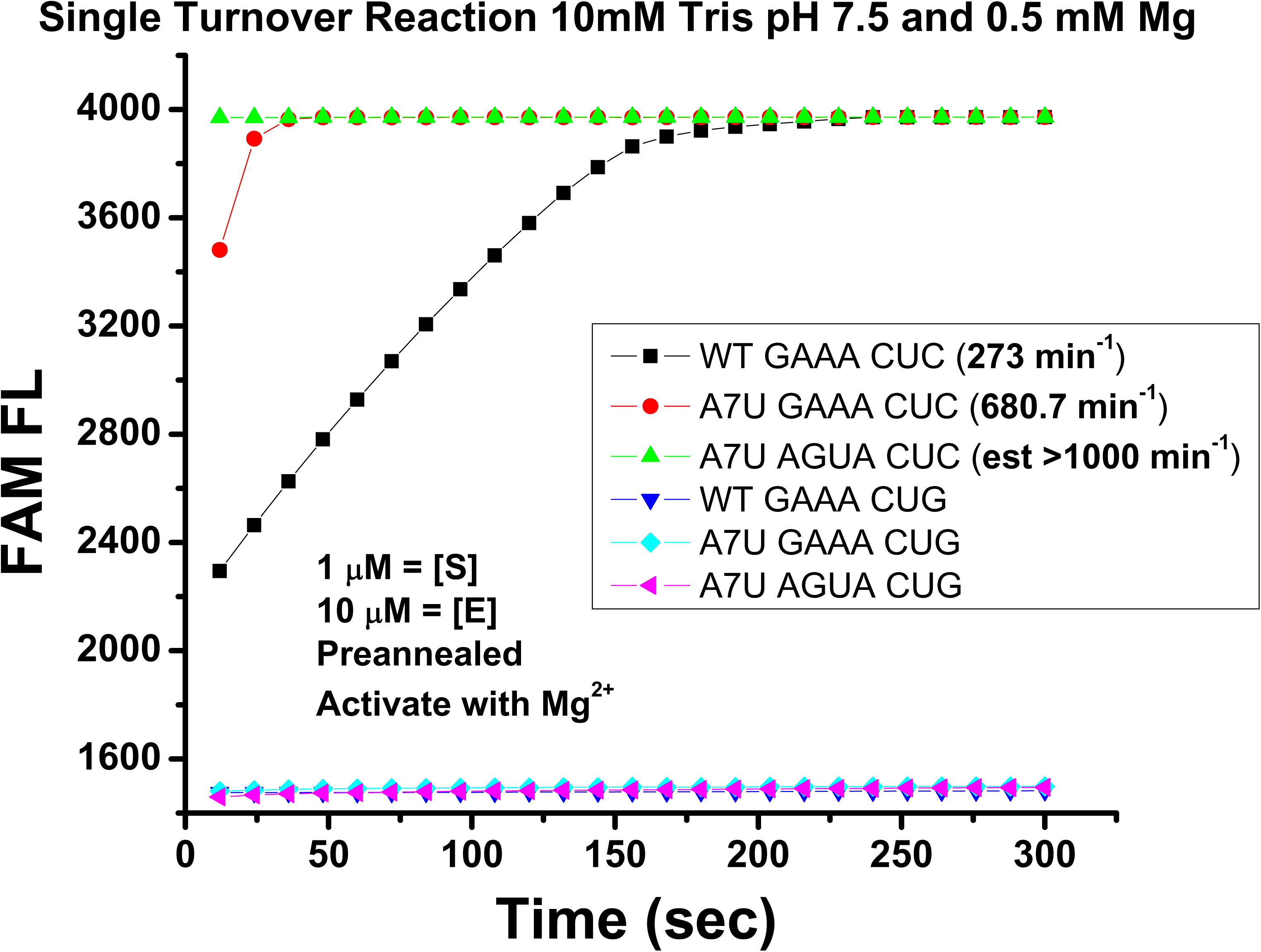
Single-turnover conditions. Optical assay in which 10 μM EhhRz was mixed with 1 μM 15-mer substrate in 10 mM Tris-HCl (pH 7.5). The solution was heated to 95°C for 2 min and 65°C for 2 min and then cooled to room temperature. Reaction was catalyzed by adding Mg^2+^ to final concentration of 0.5 mM, and reaction tubes were placed immediately (within 30 s) in the real-time PCR machine. WT(GAAA) EhhRz showed a measurable nonhyperbolic (linear) rate under these conditions; only a few points were measurable for the A7U(GAAA) to obtain an estimated rate, and the A7U(AGUA) enzyme fully cleaved the substrate by the first time point. Noncleavable substrates with CUG motifs showed lower levels of initial fluorescence and no evidence of cleavage. Catalytically inactivated EhhRzs (G8C) showed low levels of fluorescence that did not increase significantly over time (data not shown).

## DISCUSSION

The hhRz is a site-specific RNA endonuclease that was initially anticipated to have great therapeutic potential (Uhlenbeck, 1987). Ribozyme Pharmaceuticals Inc. developed systemically delivered hhRzs (Heptazyme™ targeting the 5’ end of hepatitis C virus RNA, Herzyme™ for *HER2* overexpression in breast and ovarian cancers, and Angiozyme® targeting VEGF receptor-1 for stage IV metastatic breast cancer) that were tested in clinical trials but failed to progress to clinical approval (Usman et al., 1996; Usman and Blatt, 2000; Zhou et al., 2019). In retrospect, this is not surprising given the bottlenecks in the development and delivery of posttranscriptional gene silencing agents (Sullivan et al., 2008, 2012; Juliano 2016). An injected nuclease-resistant hhRz must be distributed pharmacodynamically in serum and enter the appropriate cells in target tissues at concentrations sufficient to ensure annealing with structured target mRNA at an accessible site to exert cleavage and therapeutic knockdown. Such a multivariate problem severely constrains the probability of success with systemic delivery.

Vector-mediated gene therapy bypasses the problems associated with systemic delivery by expressing the construct in target cells (e.g., retina; Gorbatyuk et al., 2007; Cideciyian et al., 2018). Many studies, including our own, have demonstrated a range of target knockdown efficacies for hhRzs at the mRNA and protein level via gene therapy methods (Lieber and Strauss, 1995; Abdelmaksoud et al., 2009; Yau et al., 2016, 2019). With this approach, the hhRz is overexpressed relative to the target mRNA to maximize the probability of interactions at intracellular physiological Mg^2+^ levels. However, mhhRzs have a low turnover rate of <1–2 min^−1^ for small unstructured targets, which is generally orders of magnitude lower for structured full-sized mRNA targets. Notably, several computational and experimental approaches can identify rare regions of the folded native-length mRNA that are accessible to these therapeutic agents *in vitro* or *in cellulo* (Patzel and Sczakiel, 1998; Scherr and Rossi, 1998; Amarzguioui et al., 2000; Ding and Lawrence, 2001; Abdelmaksoud et al., 2009; Yau et al., 2016; Froebel et al., 2018). Improvements in hhRz kinetics, the prime variable limiting therapeutic potency, could substantially revitalize their therapeutic potential. However, when the substrate concentration exceeds that of the ribozyme (Michaelis-Menten conditions)—conditions ideal for therapeutics—the kinetics of the reaction are expected to be slower because of the lower collision frequency. Hence, substantially improved kinetics could maximize target knockdown at lower expression levels or delivery levels, and thereby lower the coincident risk for toxicity.

mhhRzs lack the TAEs found in naturally occurring “extended” hhRzs (xhhRzs) that are responsible for enhanced kinetic performance at cellular levels of free Mg^2+^ (Khvorova et al., 2003; De La Pena et al., 2003; Peneda et al., 2004). Subsequent structural analyses of xhhRz RNAs with upstream TAEs explained extensive biochemistry data that crystal structures of the mhhRz were unable to explain (Blount and Uhlenbeck, 2005; Martick and Scott, 2006; Nelson and Uhlenbeck, 2006, 2008; Scott, 2007). Stem-loop II interactions with *in cis* TAEs are not conserved and exhibit variability in nature (Shepontinovskaya et al., 2008). Although mhhRzs have a lower probability of sampling the active state structure(s), their reaction mechanisms and catalytically active structures are similar to those of xhhRzs (Nelson and Uhlenbeck, 2008). Thus, there may be many structural paths (dynamic and topological) and interactions that lead to a functional catalytically active state(s). The free energy, topology, and dynamics of a more remote loop impact the probability of a successful catalytic event (McDowell et al., 2010; Bevilacqua et al. 2019), because loop interactions initiate structural/dynamic transitions that propagate into the enzymatic core to affect chemical mechanistic processes (e.g., sn2 alignment, general base, general acid). Regardless, studies of xhhRzs have shown that this small catalytic RNA has greater kinetic capacity than initially estimated.

To enhance the therapeutic potential of hhRzs, their capacity to cleave targets *in trans* has been investigated (Saksmerprome et al., 2004). Cleavage rates of xhhRzs are generally improved versus mhhRzs under physiological or supraphysiological Mg^2+^ levels. However, to obtain a bulge loop structure in the upstream region to simulate an xhhRz functioning *in cis*, the stem I antisense flank of the hhRz *in trans* must be extended. From a therapeutic perspective, this would lower specificity (because an improperly bound target is not displaced and could still cleave) and decrease turnover (lower leaving rate of the upstream flank) (Stage-Zimmermann and Uhlenbeck, 1998; Herschlag, 1991). Furthermore, extension of stem II to generate additional nonterminal loop structures will inhibit the activity of the hhRz *in trans* (Homann et al., 1994; Clouet-d’Orval and Uhlenbeck, 1997; Yau et al., 2016, 2019). In summary, it is difficult to transition the enzymatic properties in the naturally occurring *in cis* functionality (cleavage of a target after intramolecular recognition and ligation in the rolling circle replication model) to *in trans* functionality (therapeutic intermolecular attack). Therefore, we were intrigued by the discovery of O’Rourke et al. (2015) that a highly active *in trans* mhhRz functionality (estimated at 61 min^−1^ at physiological pH in 10 mM Mg^2+^) is accomplished by including an unpaired U7 residue on the downstream substrate flank.

The loop U7 residue in the substrate component was originally identified in many but not all xhhRzs (Khvorova et al., 2003; Shepotinovskaya and Uhlenbeck, 2008), in which a specific Hoogsteen base face interaction was noted with the terminal A residue (e.g., LA4) in the GUGA (GRNA) stem II loop. The Hoogsteen LA4:U7 interaction was identified in two full-length hhRz crystal structures of the xhhRzs of *Schistosoma mansoni* (PDB 3ZP8, class I xhhRz) and the satellite RNA of tobacco ringspot virus (TRSV) (PDB 2QUS, class III xhhRz) (Anderson M et al., 2013; Chi et al., 2008). The interaction between the upstream antisense flank region and the stem II loop is critical for meaningful cleavage at cellular levels of free Mg^2+^ (≤1 mM), which is approximately an order of magnitude lower than the standard conditions used to evaluate hhRz kinetics (≥10 mM). Single-turnover cleavage rates of xhhRzs with upstream components can be 100-fold better than those of mhhRzs. Khvorova et al. (2003) measured an intramolecular (*in cis*) cleavage rate of 1.2 min^−1^ for the TRSV xhhRz at 0.1 mM Mg^2+^, pH 7, and 37°C; nine other xhhRzs have rates from <0.01 to 1.40 min^−1^. Site-specific mutagenesis revealed an interaction between the GNRA (GUGA) tetraloop capping stem II and the heptaloop (UGUGCUU) capping stem I; three of the four residues of the tetraloop were essential to cleavage activity (L2.1, L2.3, and L2.4; not L2.2 [U]) and five of the seven nucleotides in the heptaloop were essential (L1.1–L1.4 and L1.6); L1.1 is equivalent to the U7 substrate residue in stem I described by O’Rourke et al., (2015). Khvorova et al. (2003) structurally modeled several *cis* and *trans* Watson-Crick and sugar:sugar interactions between the nucleotides in the stem II and stem I loops of TRSV xhhRz (PDB 1P66). O’Rourke et al. (2015) used gel-based cleavage assays and measured a rate of 0.77 min^−1^ for their mhhRz (HH_min-AL4:U1.7_) at pH 5.6 (to slow activity) and 27°C with the mhhRz pre-annealed in 50-fold molar excess relative to its 21-mer substrate (5’ labeled with Cy3) in a reaction initiated by 10 mM Mg^2+^; rates extrapolated to pH 7.5 were 61 ± 9 min^−1^ with the assumption that pH affects only the cleavage rate (*k_2_*) and not the product leaving rate. An additional construct extended the extrapolated single-turnover cleavage rate to 95 ± 20 min^−1^ at 10 mM Mg^2+^. Control reactions in which the upstream antisense flank was extended and a U7 residue was included in a Watson-Crick base pair had lower rates than those with mhhRzs.

Single-turnover conditions in conventional gel-based cleavage assays reflect the cleavage rate of the hhRz (*k_2_*). Our novel optical turnover assay requires both the cleavage rate and at least one product dissociation rate (to separate the FAM on the upstream cleavage product from the BHQ1 on the downstream product). Under our conditions (0.25 mM Mg^2+^), which were closest to those used by Khvorova et al. (2003), the maximum scaled turnover activity was 69.70 ± 9.27 nM min^−1^ for our best EhhRz (A7U AGUA), which is 50-fold better than their maximum single-turnover rate of 1.4 min^−1^ at 0.1 mM Mg^2+^. With 10 mM Mg^2+^, simulating the conditions used by O’Rourke et al. (2015), our most active enzyme [A7U(AGUA)] had a scaled turnover rate of 486.51 ± 43.47 nM min^−1^, which is 5.1-fold better than their maximum (estimated) single-turnover activity of 95 min^−1^ assessed at 10 mM Mg^2+^. Our single-turnover rates at physiological Mg^2+^ (0.5 mM) were greater than 680 nM min^−1^ for the A7U(GAAA) EhhRz (7.16-fold) and were likely greater than 1,000 nM min^−1^ (>10.5-fold) for our best construct [A7U(AGUA)] relative to the O’Rourke et al. (2015) best construct measured under single-turnover conditions with 10 mM Mg^2+^. The measured turnover rate for HH16 (∼9 nM min^−1^) in our optical assay at 0.5 mM Mg^2+^ and 37°C (pH 7.5) is greater than the 1–2 min^−1^ measured under single-turnover conditions in various studies under “standard” conditions (25°C, 10 mM Mg^2+^, and pH 7.5) in gel based assays; this can attributed, at least in part, to the higher reaction temperature. Relative to the classic HH16, under the same substrate excess (turnover) conditions at 37°C and 0.5 mM Mg^2+^ the improvement is 18.8-fold for our A7U(AGUA) enzyme with our assay. Our best EhhRz has the same enzyme core and the same stem II sequence as HH16; the only differences are between the stem II tetraloop and the antisense flanks.

There are other reports in the literature of enhanced hhRz activity. The gel-based assays supporting these observations are generally conducted under non-Michaelis-Menten conditions of enzyme excess [pre-annealed [E]>>[S]) and nonphysiological “standard” Mg^2+^ conditions (≥10 mM Mg^2+^). Clouet-d’Orval and Uhlenbeck (1997) identified a class I/II mhhRz (HH1α) with single-turnover activity of 10 min^−1^ under standard conditions. By grafting elements of HH1α onto HH16 (0.4 min^−1^) and decreasing the pH to 6.5 to slow kinetics, they determined that the first two base pairs (U:A/A:U) of the stem I antisense flank were responsible for enhanced activity and that a stem II length >4 bp was inhibitory. Canny et al. (2004) reported very high cleavage rates (∼870 min^−1^, 25°C) from an *in trans* form of the *Schistosoma* xhhRz tested under single-turnover (enzyme excess) conditions with very high levels of Mg^2+^ (200 mM; *K_d_*, ∼40 mM) and high pH (8.5), which, though far from physiological, demonstrated the *catalytic potential* of RNA enzymes; U6 and U7 residues were present in the substrate strand, and the stem I antisense flank had a total of 12 nt. Saksmerprome et al. (2004) used evolutionary techniques to identify an *in trans* hhRz (“RzB”) which simulated the structure and function of the peach latent mosaic viroid xhhRz operating *in cis.* RzB uses a long stem I antisense flank (11 nt) with a central UAA bulge that interacts with a 6-nt stem II loop (5’-UGGGAU-3’, similar to the naturally occurring sequence of peach latent mosaic viroid hhRz [UGAGAU]). RzB demonstrated rates of 1.8 min^−1^ at 37°C with 0.5 mM Mg^2+^ under single-turnover pre-annealed reaction conditions with enzyme in 10-fold molar excess over substrate, which did not harbor a U7 residue. Roychowdhury and Burke (2006) investigated pre-steady state reaction conditions with the *in trans* RzB and measured “extraordinary” cleavage rates (780 min^−1^ at pH 7.4, 37°C, and 1 mM Mg^2+^) in a stopped-flow apparatus with substrate and enzyme pre-annealed in complex (10:1 [E]/[S]).

Reactions occurred with multiphasic kinetics, and the fastest single-turnover rate component represented only ∼5% of the overall cleavage extent, which otherwise progressed to 80% cleavage. This study established the upper limit of the hhRz cleavage kinetics at the time and suggested means to achieve such efficiencies. The goal was to utilize RzB for intracellular target knockdown applications (Saksmerprome et al., 2004), and it was initially tested *in vitro* against a strongly structured fragment of HIV RNA, for which cleavage was orders of magnitude slower. We performed a translational study with the RzB format, embedded internally within a modified VA1 scaffold RNA (Lieber and Strauss, 1995; Prislei et al., 1997), but it did not improve target knockdown relative to that with an mhhRz without TAEs targeting the same accessible site (Yau et al., 2019).

### Practical implications for clinical or biotechnological translation

Our studies clearly show that the EhhRz A7U(AGUA) achieves scaled turnover rates of 314 nM min^−1^ under steady-state (substrate excess) conditions at cellular levels of free Mg^2+^ (≤1 mM). This EhhRz achieves saturation cleavage of 1 μM substrate within 5 min, as expected for this level of turnover at 0.5 mM Mg^2+^ (**Fig. 1B**). Our EhhRz is designed to cleave an accessible region in *RHO* mRNA, a target for knockdown/reconstitute therapy in autosomal dominant retinitis pigmentosa and knockdown therapy in age-related macular degeneration (Sullivan et al, 2002; Abdelmaksoud et al., 2009; Yau et al., 2019). The efficiency of our fastest enzyme to date [A7U(AGUA)] (V_max_ = 214.62 ± 27.92 min^-1^, K_m_ = 1344.08 ± 211.13 nM) is 1.60 × 10^8^ M^−1^min^−1^, which is over 1 log-fold more efficient than the 525 *RHO* mhhRz (Gorbatyuk et al., 2007), which had a *K_m_* of 152 nM and *k*_cat_ of 0.78 min^−1^ under substrate-excess conditions at 2 mM Mg^2+^ and 37°C, or an efficiency of 5.13 × 10^6^ M^−1^ min^−1^. The *K_m_* of the EhhRz tested here is on the same scale, but the *k*_obs_ at lower levels of Mg^2+^ (0.5 mM) is over two orders of magnitude faster, making for an enzymatic efficiency that rivals that of RNase A for 266 A7U(AGUA). The *K_m_* of the enzyme is largely related to the antisense flank energy required for molecular recognition (binding) of the target sequence and will predictably vary with antisense flank length and composition, which can be optimized to maximize selectivity and product leaving rates (Stage-Zimmermann and Uhlenbeck, 1998). The *k*_obs_ is a major chemical property of the enzyme proper, for which EhhRzs clearly demonstrate log-order improvement over hhRzs or xhhRzs. When evaluating EhhRzs under single-turnover conditions (10-fold molar excess of enzyme over substrate [at 1 μM]), pre-annealed and initiated by 0.5 mM Mg^2+^, we were unable to measure the rate with current optical methods of the A7U(AGUA) enzyme because of the rapid speed (estimated >1000 nM min^-1^), we can only estimate the A7U(GAAA) enzyme at >680 nM min^−1^ and obtain a rate of approximately 273 nM min^−1^ for the WT(GAAA) enzyme. We did not explore reduced pH, because the optical assay designed here depends upon product release, and base pairing can be affected by pH (Thapylal et al., 2014; Szabat and Kierzek, 2017; O’Connell et al., 2019). Rapid kinetic assays (stopped-flow) are needed to investigate these rates under physiological conditions (Bingaman et al., 2017).

What are the reasons for this exceptional catalytic performance? EhhRz 266 A7U(AGUA) achieves rates well beyond the expected performance of an mhhRz. Adopting the perspectives of Emilsson and Breaker (2003), Breaker et al. (2003), Cochrane and Strobel (2008), and Bevilacqua et al. (2019), a logical explanation is that this EhhRz uses more of the available mechanisms for rate enhancement than just the α (sn2-orientational alignment of the 2’-OH, the scissile bond, and the proton donor) and γ (general base mechanism to abstract a proton from the 2’-OH, thought to relate to properties of G12) processes used by the mhhRz. The additional use of the β (decreasing the charge accumulation on the phosphate of the activated state) and δ (efficient protonation of the leaving group by the general acid, thought to relate to properties of G8) mechanisms could bring the catalytic efficiency of the EhhRz to the scale of RNase A, which is thought to use all four physical mechanisms for phosphodiester cleavage.

The GNRA tetraloops consistently supported high turnover activity for the model EhhRz, whereas the single model UUCG and CUUG tetraloops poorly supported high turnover activity. The first and last nucleotides of these canonical tetraloops form base pairs, and these tetraloops have varied stability (Thapar et al., 2013; Klosterman et al., 2004; Dale et al., 2000; Molinaro and Tinoco, 1995; Tuerk et al., 1988). These tetraloops also have varied structures, which do not shed light on the enhanced activity of the EhhRzs studied here (Bottaro and Lindorf-Larsen, 2017). The distinct empirical finding is in the A7U(AGUA) construct, which uses a less-well-studied AGNN tetraloop that does not have an L1:L4 base pair. Rather, the first and second nucleotides stack on the underlying 5’ nucleotide of the closing base pair of the underlying stem (stem II in hhRz) (Wu et al., 2001, 2004; Thapar et al., 2014) and the second G residue is in a *syn* configuration, which causes outpocketing of the phosphodiester bond with the third nonconserved N residue (Lebars et al., 2001). It is the A7U(AGUA) EhhRz construct that uniquely demonstrates the most rapid kinetics and a clear cooperative transition in the presence of Mg^2+^. The roles of different tetraloops have evolved for interactions with other RNAs or proteins. One may speculate that the unique structural and thermodynamic properties of the AGUA tetraloop (and, to a similar extent, the AUUA, AAUA, and AAAA tetraloops; see **Fig. 3A, B**) facilitate the Mg^2+^-dependent conformational changes that drive and stabilize a highly efficient catalytically active state that maximizes the engagement of all four potential contributors (α, γ, β, and δ) to accelerate phosphodiester cleavage. Biophysical, molecular dynamics, and structural biological experiments are needed to better understand the enhancements that we have observed.

There are several consequences of our observations. *First,* an EhhRz with such substantial improvements in enzymatic performance under physiological conditions requires lower levels of expression in/delivery to target cells to achieve therapeutic target mRNA knockdown, thus reducing the toxicity from RNA-sensing pathways (e.g., RIG, PKR) or off-target effects, which are concentration dependent. EhhRzs thus revitalize the potential of catalytic nucleic acid therapeutics. *Second*, the enzymatic potential of the hhRz is not yet fully explored. The broad range of rates from a limited variety of stem II loop cap mutants strongly suggests that the structural and functional mechanistic biology of EhhRzs have further catalytic potential that incentivizes further investigation. It also shows that RNA catalytics with native chemistry (G, U, C, and A) have the capacity for turnover rates that are much better than anticipated for the mhhRz format at 1–2 min^−1^, which utilizes only two of four rate acceleration mechanisms (Emilsson et al., 2003; Breaker et al., 2003). *Third,* given that all the stem II loop variants had identical antisense flanks and that the reaction and buffer conditions were otherwise identical, the product leaving rates should also be identical. As a result, the cleavage rate, *k2*, cannot be a constant as previously inferred (Stage-Zimmermann and Uhlenbeck, 1998). *Fourth,* because the stem II capping loop can be functionally varied to enhance turnover rates with variable stem I interactions, the loop sequence could be optimized for particular upstream flanks (receptors) in arbitrary targets (consider GNRA tetraloop receptors; Abramovitz et al., 1997; Geary et al., 2008; Fiore and Nesbitt, 2013). *Fifth,* the small size of the EhhRz inspires investigations of novel synthetic catalytics with non-natural nucleotides at specific positions for further rate enhancement. The enzyme efficiency (*k*_cat_/*K_m_*), which already rivals that of RNase A, might be improved further if the *k*_cat_ can be improved by nucleotide chemistries tuned to enhance the known mechanism of reaction (e.g., designing the pK_a_s of G12 and G8 residues to bring them closer to neutrality, as in RNase A, which uses two histidine residues [general base H12 and general acid H119] in the active site) (Thompson et al., 1995; Emilsson et al., 2003; Breaker et al., 2003; Han and Burke, 2005; Strobel and Cochrane, 2007; Cochrane and Strobel, 2008). *Sixth,* recently, it was shown that synthetic antisense oligodeoxynucleotides injected into the vitreous of the eye (a closed volume space) can be used to treat mutant pre-mRNA splicing defects in photoreceptor nuclei (Cideciyan et al., 2019); this study speaks volumes to a new emerging class of nucleic acid therapeutics for outer retinal degenerative conditions, or perhaps ocular conditions in general. *Seventh,* based on a prior preclinical study (Murray et al., 2015) an ongoing human clinical trial (NCT04123626; clinicaltrials.gov; ProQR Therapeutics, QR-1123) is evaluating use of next generation antisense agents, injected into the human vitreous body, in order to suppress only the mutant P23H rhodopsin mRNA of patients with an autosomal dominant form of retinitis pigmentosa, an inherited retinal dystrophy; use of EhhRzs in this context could provide a substantial improvement to antisense agents which have slower kinetic turnover and must usurp host cell machinery (e.g., RNasH) for cleavage of target mRNAs. As we begin to explore the biophysics and structural biology of EhhRzs, it is already clear that catalytic capacity is not yet fully understood and likely to have greater potential. These newly described properties of EhhRzs revitalize the potential for their use in human nucleic acid therapeutics or in a biotechnological capacity (e.g., aptazymes). We expect that our work could open a new phase of advanced biophysical, chemical, structural biological, and evolutionary studies on the EhhRz as a nucleic acid enzyme and as a therapeutic.

## DATA AVAILABILITY

Datasets for the Figures in the manuscript are deposited with Dryad (Citation number to be obtained prior to submission to Nucleic Acids Research).

## REFERENCES

Abdel-Maksoud H, Yau EH, Zuker M, Sullivan JM. Development of lead hammerhead ribozyme candidates against human rod opsin for retinal degeneration therapy. Experimental Eye Research 2009; 88(5): 859–879.

Abramovitz DL, Pyle AM. Remarkable morphological variability of a common RNA folding motif: The GNRA tetraloop-receptor interaction. J. Mol. Biol. 1997; 266:493–506.

Altman S. Ribonuclease P. J. Biol. Chem. 1990; 265: 20053–20056.

Amarzguioui M, Brede G, Babale E, Grotli M, Sproat B, Prydz H. Secondary structure prediction and *in vitro* accessibility of mRNA as tools in the selection of target sites for ribozymes. Nucleic Acids Res. 2000; 28:4113–4124

Anderson M, Schultz EP, Martick M, Scott WG. Active-site monovalent cations revealed in a 1.55-Å-resolution hammerhead ribozyme structure. J. Mol. Biol. 2013; 425:3790–3798.

Antczak M, Popenda M, Zok T, Sarzynska J, Ratajczak T, Tomczyk K, Adamiak RW, Szachniuk M. New functionality of RNAComposer: an application to shape the axis of miR160 precursor structure. Acta Biochimica Polonica 2016; 63(4): 737–744.

Bassi GS, Murchie AI, Lilley DM. The ion-induced folding of the hammerhead ribozyme: core sequence changes that perturb folding into the active conformation. RNA 1996 Aug; 2(8): 756–768.

Bassi GS, Murchie AI, Walter F, Clegg RM, Lilley DM. Ion-induced folding of the hammerhead ribozyme: a fluorescence resonance energy transfer study. EMBO J 1997 Dec 15; 16(24):7481–7489.

Bevilacqua PC, Harris ME, Piccirilli JA, Gaines C, Ganguly A, Kostenbader K, Ekesan S, York DM. An ontology for facilitating discussion of catalytic strategies of RNA-cleaving enzymes. ACS Chem. Biol. 2019 June 21; 14(6): 1068–1076.

Bingaman JL, Messina KJ, Bevilacqua PC. Probing fast ribozyme reactions under biological conditions with rapid quench-flow kinetics. Methods 2017 May 1; 120:125–134.

Birikh KR, Heaton PA, Eckstein F. The structure, function and application of the hammerhead ribozyme. Eur. J. Biochem. 1997; 245: 1–16.

Blount KF, Uhlenbeck OC. The structure-function dilemma of the hammerhead ribozyme. Annu. Rev. Biophys. Biomol. Struct. 2005: 34: 415–440.

Bottaro S, Lindorff-Larsen K. Mapping the universe of RNA tetraloop folds. Biophys. J. 2017; 113:257–267.

Breaker RR, Emilsson GM, Lazarev D, Nakamura S, Puskarz IJ, Roth A, Sudarsan N. A common speed limit for RNA-cleaving ribozymes and deoxyribozymes. RNA 2003; 9: 949–957.

Busan S, Weidmann CA, Sengupta A, and Weeks KM. Guidelines for SHAPE reagent choice and detection strategy for RNA structure probing studies. Biochemistry 2019; 58: 2655–2664.

Canny MD, Jucker FM, Kellogg E, Khorova A, Jayasena SD, et al., Fast cleavage kinetics of a natural hammerhead ribozyme. J. Am. Chem. Soc. 2004; 126: 10848–10849.

Cech TR. Self-splicing of group I introns. Annu. Rev. Biochem. 1990; 59: 543–568.

Chi Y-I, Martick M, Lares M, Kim R, Scott WG, Kim S-H. Capturing hammerhead ribozyme structures in action by modulating general base catalysis. PLoS Biol. 2008; 6:e234.

Cideciyian AV, Sudharsan R, Dufour VL, Massengill MT et al., Mutation-independent rhodopsin gene therapy by knockdown and replacement with a single AAV vector. Proc. Natl. Acad. Sci. USA 2018 Sep 4; 115(36): E8547–8556.

Cideciyan AV, Jacobson SG, Drack AV, Ho AC et al., Effect of an intravitreal antisense oligonucleotide on vision in Leber congenital amaurosis due to a photoreceptor cilium defect. Nat. Med. 2019 Feb; 25(2):225–228.

Clouet-d’Orval B, Uhlenbeck OC. Hammerhead ribozymes with a faster cleavage rate. Biochemistry 1997; 36: 9087–9092.

Cochrane JC and Strobel SA. Catalytic strategies of self-cleaving ribozymes. Acc. Chem. Res. 2008; 41(8): 1027–1035.

Conrad T, Plumbom I, Alcobendas M, Vidal R, Sauer S. Maximizing transcription of nucleic acids with efficient T7 promoters. Communications Biol. 2020; 3:439.

Dahm SAC, Derrick WB, Uhlenbeck OC. Evidence for the role of solvated metal hydroxide in the hammerhead cleavage mechanism. Biochemistry 1993; 32:13040–13045.

Dale T, Smith R, Serra MJ. A test of the model to predict unusually stable RNA hairpin loop stability. RNA 2000; 6: 608–615.

De la Peña M, Gago S, Flores R. Peripheral regions of natural hammerhead ribozymes greatly increase their self-cleavage activity. EMBO 2003; 22: 5561–5570.

Ding Y, Lawrence CE. Statistical prediction of single-stranded regions in RNA secondary structure and application to predicting effective antisense target sites and beyond. Nucleic Acids Res. 2001; 29:1034–1046

Dufour D, del la Peña, Gago S, Flores R, Gallego J. Structure-function analysis of the ribozymes of chrysanthemum chlorotic mottle viroid: a loop-loop interaction motif conserved in most natural hammerheads. Nucleic Acids. Res. 2009; 37(1): 368–381.

Emilsson GM, Nakamura S, Roth A, Breaker RR. Ribozyme speed limits. RNA 2003; 9: 907–918.

Fiore JL, Nesbitt DJ. An RNA folding motif: GNRA tetraloop-receptor interactions. Quart. Revs. Biophys. 2013; 46(3):223–264.

Gaudin C, Ghazal G, Yoshizawa S, Abou Elela S, Fourmy D. Structure of an AAGU tetraloop and its contribution to substrate selection by yeast RNase III. J. Mol. Bio. 2006; 363: 322–331.

Geary C, Baudrey S, Jaeger L. Comprehensive features of natural and *in vitro* selected GNRA tetraloop-binding receptors. Nucleic Acids Res. 2008; 36(4): 1138–1152.

Gorbatyuk M, Justilien V, Liu J, Hauswirth WW, Lewin AS. Preservation of photoreceptor morphology and function in P23H rats using an allele independent ribozyme. Exp. Eye Res. 2007 Jan; 84(1): 44–52.

Hammann C, Norman DG, Lilley MJ. Dissection of the ion-induced folding of the hammerhead ribozyme using ^19^F NMR. Proc. Natl. Acad. Sci. USA 2001 May 8; 98(10): 5503–5508.

Han J, Burke JM. Model for general acid-base catalysis by the hammerhead ribozyme: pH activity relationships of G8 and G12 variants in the putative active site. Biochemistry 2005; 44:7864–7870.

Haseloff J, Gerlach WL. Simple RNA enzymes with new and highly specific endoribonuclease activities. Nature 1988 August; 334:585–591.

Hertel KJ, Pardi A, Uhlenbeck OC, Koizumi M, Ohtsuka E, Uesugi S., et. al. Numbering system for the hammerhead. Nucleic Acids Res. 1992; 20:3252.

Hertel KJ, Herschlag D, Uhlenbeck OC. A kinetic and thermodynamic framework for the hammerhead ribozyme reaction. Biochemistry 1994; 33: 3374–3385.

Hertel KJ, Uhlenbeck OC. The internal equilibrium of the hammerhead ribozyme reaction. Biochemistry 1995; 34: 1744–1749.

Hertel KJ, Herschlag D, Uhlenbeck OC. Specificity of hammerhead ribozyme cleavage. EMBO J. 1996; 15(14):3751–3757.

Hertel KJ, Peracchi A, Uhlenbeck OC, Herschlag D. Use of intrinsic binding energy for catalysis by an RNA enzyme. Proc. Natl. Acad. Sci. USA 1997; 94: 8497–8502.

Homann M, Tabler M, Tzortzakaki S, Sczakiel G. Extension of helix II of an HIV-1-directed hammerhead ribozyme with long antisense flanks does not alter kinetic parameters in vitro but causes loss of the inhibitory potential in living cells. Nucleic Acids. Res. 1994 Sep 25; 22(19):3951–3957.

Juliano RL. The delivery of therapeutic oligonucleotides. Nucleic Acids Res. 2016 Aug 19; 44(14): 6518–6548.

Khvorova A, Lescoute A, Westhof E, Jayasena SD. Sequence elements outside the hammerhead ribozyme catalytic core enable intracellular activity. Nat. Struct. Biol. 2003; 10: 708–712.

Kim N-K, Murali A, DeRose VJ. Separate metal requirements for loop interactions and catalysis in the extended hammerhead ribozyme. J. Amer. Chem. Soc. 2005; 127:14134–14135.

Klosterman PS, Hendrix DK, Tamura M, Holbrook SR, Brenner SE. Three-dimensional motifs from the SCOR, structural classification of RNA database, extruded strands, base triple, tetraloops and U-turns. Nucleic Acids Res. 2004; 32(8): 2342–2352.

Lieber A, Strauss M. Selection of efficient cleavage sites in target RNAs by using a ribozyme expression library. Mol. Cell. Biol. 1995; 15: 540–551

Lilley DMJ. Ribozymes—a snip too far? Nat. Struct. Biol. 2003 Sep; 10(9):672–3.

Martick M, Scott WG. Tertiary contacts distant from the active site prime ribozyme for catalysis. Cell 2006; 126:309–320.

McDowell SE, Jun JM, Walter NG. Long-range tertiary interactions in single hammerhead ribozymes bias motional sampling toward catalytically active conformations. RNA 2010; 16:2414–2426.

Menger M, Tuschl T, Eckstein F, Porschke D. Mg^2+^-dependent conformational changes in the hammerhead ribozyme. Biochemistry 1996; 35:14710–14716.

Menger M, Eckstein F, and Porschke D. Multiple conformational states of the hammerhead ribozyme, broad time range of relaxation and topology of dynamics. Nucleic Acids Res. 2000 Nov 15; 28(22): 4428–4434.

Mir A, Chen J, Robinson K, Lendy E, Goodman J, Neau D, Golden BL. Two divalent metal ions and conformational changes play roles in the hammerhead ribozyme cleavage reaction. Biochemistry 2015 Oct 20; 54(41): 6369–6381.

Mir A, Golden BL. 2016. Two active site divalent ions in the crystal structure of the hammerhead ribozyme bound to a transition state analogue. Biochemistry 2016 Feb 2; 55(4):633–636.

Murray SF, Jazayeri A, Matthes MT, Yasumura D, Yang H, Peralta R, Watt A, Freier S, Hung G, Adamson PS, Guo S, Monia BP, LaVail MM, McCaleb ML. Allele-specific inhibition of rhodopsin with an antisense oligonucleotide slows photoreceptor degeneration. Invest. Ophthalmol. Vis. Sci. 2015 Oct; 56(11): 6362–6375.

Myers JM, Sullivan JM. RNA Structure-Function Properties in Facilitated Hammerhead Ribozymes with Unprecedented Catalytic Rates. IOVS 2020 June, 61: 2293.

Nelson JA, Uhlenbeck OC. When to believe what you see. *Molec*. Cell 2006; 23(4):447–450.

Nelson JA, Uhlenbeck OC. Minimal and extended hammerheads utilize a similar dynamic reaction mechanism for catalysis. RNA 2008; 14:43–54.

Nelson JA, Uhlenbeck OC. Hammerhead redux: does the new structure fit the old biochemical data? RNA 2008; 14: 605–615.

O’Connell AO, Hanson JA, McCaskill DC, Moore ET, Lewis DC, and Grover N. Thermodynamic examination of pH and magnesium effect on U6 RNA loop. RNA 2019; 25: 1779–1792.

O’Rourke SM, Estell W, Scott WG. Minimal hammerhead ribozymes with uncompromised catalytic activity. J. Mol. Biol. 2015; 427:2340–2347.

Patzel V, Sczakiel G. Theoretical design of antisense RNA structures substantially improves annealing kinetics and efficacy in human cells. Nat. Biotechnol. 1998; 16: 64–68.

Penedo JG, Wilson TJ, Jayasena SD, Khvorova A, Lilly DM. Folding of the natural hammerhead ribozyme is enhanced by interaction of auxiliary elements. RNA 2004; 10: 880–888.

Popenda M, Szachniuk M, Antczak M, Purzycka KJ, Lukasiak P, Barton N, Blazewicz J, Adamiak RW. Automated 3D structure composition for large RNAs. Nucleic Acids Res. 2012; 40(14): e112.

Prislei S, Buonomo SBC, Michienzi A, Bozzoni I. Use of adenoviral VAI small RNA as a carrier for cytoplasmic delivery of ribozymes. RNA 1997; 3: 677–687.

Reuter JS, Mathews DH. RNAstructure: software for RNA secondary structure prediction and analysis. BMC Bioinformatics 2010 Mar 15; 11:129.

Roychowdhury-Saha M, Burke DH. Extraordinary rates of transition metal ion-mediated ribozyme catalysis. RNA 2006; 12: 1846–1852.

Rueda D, Wick K, McDowell SE, Walter NG. Diffusely bound Mg2+ ions slightly reorient stems I and II of the hammerhead ribozyme to increase the probability of formation of the catalytic core. Biochemistry 2003 Aug26; 42(33):9924–9936.

Ruffner DE, Stormo GD, Uhlenbeck OC. Sequence requirements of the hammerhead RNA self-cleavage reaction. Biochemistry 1990; 29: 10695–10702.

Saksmerprome V, Roychowdhury-Saha M, Jayasena S, Khvorova A, and Burke DH. Artificial tertiary motifs stabilize *trans-*cleaving hammerhead ribozymes under conditions of submillimolar divalent ions and high temperatures. RNA 2004; 10:1916–1924.

Scherr M, Rossi JJ. Rapid determination and quantitation of the accessibility to native RNAs by antisense oligodeoxynucleotides in murine cell extracts. Nucleic Acids Res. 1998; 26: 5079–5085.

Shepotinovskaya IV, Uhlenbeck OC. Catalytic diversity of extended hammerhead ribozymes. Biochemistry 2008; 47: 7034–7042.

Smola MJ and Weeks KM. In-cell RNA structure probing with SHAPE−MaP. Nature Prot. 2018; 13(6): 1181–1195.

Stage-Zimmermann TK, Uhlenbeck OC. Hammerhead ribozyme kinetics. RNA 1998; 4: 875–889.

Strobel SA, and Cochrane JC. RNA catalysis: ribozymes, ribosomes, and riboswitches. Curr. Opin. Chem. Biol. 2007 Dec; 11(6): 636–643.

Sullivan JM, Myers JM. A facilitated hammerhead ribozyme rhodopsin therapeutic with catalytic efficiency on par with ribonuclease A at cellular free Mg2+ levels. IOVS 2020 June, 61: 4481.

Sullivan JM, Yau EH, Taggart RT, Butler MC, Kolniak TA. Bottlenecks in development of therapeutic post-transcriptional gene silencing agents. Vis. Res. 2008; 48: 453–469.

Szabat M, Kierzek R. Parallel-stranded DNA and RNA duplexes-structural features and potential applications. The FEBS J. 2017; 284: 3986–3998h

Tang J, Breaker RR. Examination of the catalytic fitness of the hammerhead ribozyme by in vitro selection. RNA 1997; 3(8): 914–925.

Thapar R, Denmon AP, Nikonowicz EP. Recognition modes of RNA tetraloops and tetraloop-like motifs by RNA-binding proteins. WIREs RNA 2014; 5:49–67.

Thaplyal P and Bevilacqua PC. Experimental approaches for measuring pK_a_’s in RNA and DNA. Methods Enzymol. 2014; 549: 189–219.

Thompson JE, Kutateladze TG, Schuster MC, Venegas FD, Messmore JM, and Raines RT. Limits to catalysis by Ribonuclease A. Bioorg. Chem. 1995; 23(4): 471–481.

Trujillo AJ, Myers JM, Fayazi ZS, Butler MC, Sullivan JM. A discovery with potential to revitalize hammerhead ribozyme therapeutics for treatment of inherited retinal degenerations. Adv. Exp. Med. Biol. 2019; 1185:119–124.

Tuerk C, Gauss P, Thermes C, Groebe DR, Gayle M, Guild N, Stormo G, D’Aubenton-Carafa Y, Uhlenbeck OC, Tinoco I Jr, Brody EN, Gold L. CUUCGG hairpins: extraordinarily stable RNA secondary structures associated with various biochemical processes. Proc. Natl. Acad. Sci. USA 1988; 85: 1364–1368.

Uhlenbeck OC. A small catalytic oligoribonucleotide. Nature 1987 August; 328:596–600.

Uhlenbeck OC. Less isn’t always more. RNA 2003; 9:1415–1417.

Usman N, Beigelman L, McSwiggen JA. Hammerhead ribozyme engineering. Curr. Opin. Struct. Biol. 1996; 6(4): 527–533.

Usman N, Blatt LM. Nuclease-resistant synthetic ribozymes: developing a new class of therapeutics. J. Clin. Invest. 2000; 106(10): 1197–1202.

Vendeix FAP, Munoz AM, Agris PF. Free energy calculation of modified base-pair formation in explicit solvent: a predictive model. RNA 2009; 15(12): 2278–2287.

Venditti V, Niccolai N, Butcher SE. Measuring the dynamic surface accessibility of RNA with the small paramagnetic molecule TEMPOL. Nucleic Acids Res. 2008; 36(4): e20.

Wu H, Yang PK, Butcher SE, Kang S, Chanfreau G, Feigon J. A novel famly of RNA tetraloop structure forms the recognition site for *Saccharomyces cerevisiae* RNase III. EMBO J. 2001; 20(24): 7240–7249.

Wu H, Henras A, Chanfreau G, Feigon J. Structural basis for recognition of the AGNN tetraloop RNA fold by the double-stranded RNA-binding domain of Rnt1p RNase III. Proc. Natl. Acad. Sci. USA 2004; 101(22): 8307–8312.

Yau EH, Butler MC, and Sullivan JM. A cellular high-throughput screening approach for therapeutic trans-cleaving ribozymes and RNAi against arbitrary mRNA disease targets. Exp. Eye Res. 2016 Oct; 151: 236–255.

Yau EH, Taggart RT, Zuber M, Trujillo AJ, Fayazi ZS, Butler MC, Sheflin LG, Breen JB, Yu D, Sullivan JM. Systematic Screening, Rational Development, and Initial Optimization of Efficacious RNA Silencing Agents for Human Rod Opsin Therapeutics. Transl Vis Sci Technol. 2019 Dec 12;8(6):28.

Zamel R, Poon A, Jaikaran D, Andersen A, Olive J, De Abreu D, Collins RA. Exceptionally fast self-cleavage by a *Neurospora* Varkud satellite ribozyme. Proc. Natl. Acad. Sci. USA 2004 Feb10; 101(6): 1467–1472.

Zhou L-Y, Qin Z, Zhu Y-H, He Z-Y, and Xu T. Current RNA-based therapeutics in clinical trials. Curr. Gene Ther. 2019; 19: 172–196.

Zuker M. Mfold web server for nucleic acid folding and hybridization prediction. Nucleic Acids. Res. 2003 Jul 1; 31(13): 3406–3415.

