## Supplemental Legends Figures Table for "Log-order improved *in trans* hammerhead ribozyme turnover rates: reevaluating therapeutic space for small catalytic RNAs"

#### Supplementary Materials

#### Figure Legends

**Supp. Fig. 1**. 266 hRHO Substrate and Product Fluorescence-Concentration Relationships. (**Top panel**) Relationship of 15-mer 266 Substrate RNA concentration to its fluorescence. The substrate has a 5' FAM fluor and a 3' BHQ1 quencher. (**A**) The concentration was varied from 50 nM to 5000 nM and fluorescence measured over a 5 minute period. Fluorescence emissions were stable over time at all concentrations. (**B**) The relationship of fluorescence to concentration was well-fit by a linear relationship ($R^2$ = 0.99602) with a slope of 0.25836 FAM units/nM. (**Middle panel**). Relationship of 14-mer 266 Substrate RNA concentration to its fluorescence. The substrate has a 5' FAM fluor and a 3' BHQ1 quencher. (**C**) The concentration was varied from 50 nM to 2,500 nM and fluorescence measured over a 5 minute period. Fluorescence emissions were stable over time at all concentrations. (**D**) The relationship of fluorescence to concentration was well-fit by a linear relationship ($R^2$ = 0.99235) with a slope of 0.47175 FAM units/nM. (**Bottom Panel**). Relationship of 8-mer hRHO 266 Product RNA concentration to its fluorescence. The product RNA has a 5' FAM fluor and no BHQ1 quencher at the 3' end. The 8-mer product is the same for both the 15-mer and 14-mer substrates. The concentration was varied from 7.81 nM to 600 nM and fluorescence measured over a 5 minute period (**E**). Fluorescence emissions were stable over time at all concentrations. (**F**) The relationship of fluorescence to concentration was well-fit by a linear relationship ($R^2$ = 0.99788) with a slope of 2.63555 FAM units/nM. The results of separate experiments (n=3) to measure the 8-mer 266 product specific fluorescence found Mean $\pm$ SD at 3.11455 $\pm$ 0.435 FAM Units/nM; the scale factor of 3.11 FAM Units/nM was used to scale all quantitative fluorescence measures in these experiments.

**Supp. Fig. 2**. HH16 Product Fluorescence-Concentration Relationships. Relationship of 10-mer HH16 product RNA concentration to its fluorescence. The HH16 product has a 5' FAM fluor and no BHQ1 quencher at the 3' end. The concentration was varied from 15.625 nM to 1000 nM and fluorescence measured over a 5 minute period (**A**). Fluorescence emissions were stable over time at all concentrations. (**B**) The relationship of fluorescence to concentration was well-fit by a linear relationship ($R^2$ = 0.99125) with a slope of 0.97838 FAM units/nM.

**Supp. Fig. 3**. Substrate Structure RNA Folding Analysis for Target RNAs. RNAStructure was used to fold the substrate RNAs for hRHO 266 and HH16 into secondary structures (**A, D**). There is no predicted secondary structure for 266 hRHO substrate (here 15-mer) and predicted secondary structure for HH16 substrate (18-mer) involving 4 bps. RNA Composer was used to examine possible tertiary structures for the 266 and HH16 substrates. The hRHO 15-mer substrate yields no apparent tertiary structures (**B, C**). The HH16 substrate yields tertiary structures shows from two perspectives (**E, F**).

**Supp. Table 1**. Table shows kinetic parameters calculated for Stage-Zimmermann and Uhlenbeck (1998) HhRz Model with Outcomes for the WT(GAAA) and A7U (GAAA/AGUA) *hRHO* 266 EhhRzs for both the 15-mer and 14-mer substrates. The kinetic model diagram (box) shows the Kd values and rate constants calculated for the A7U(GAAA/AGUA) EhhRzs solely for the 15-mer substrate. The A7U(GAAA) and A7U(AGUA) EhhRzs have identical antisense flanks and so have the same calculated kinetic properties. ***Kd*** values have units of **nM**. Rate constants (***k***) have units of **$min^{-1}$**.

Supp. Fig 1

### Supplementary Figure 1, Myers and Sullivan

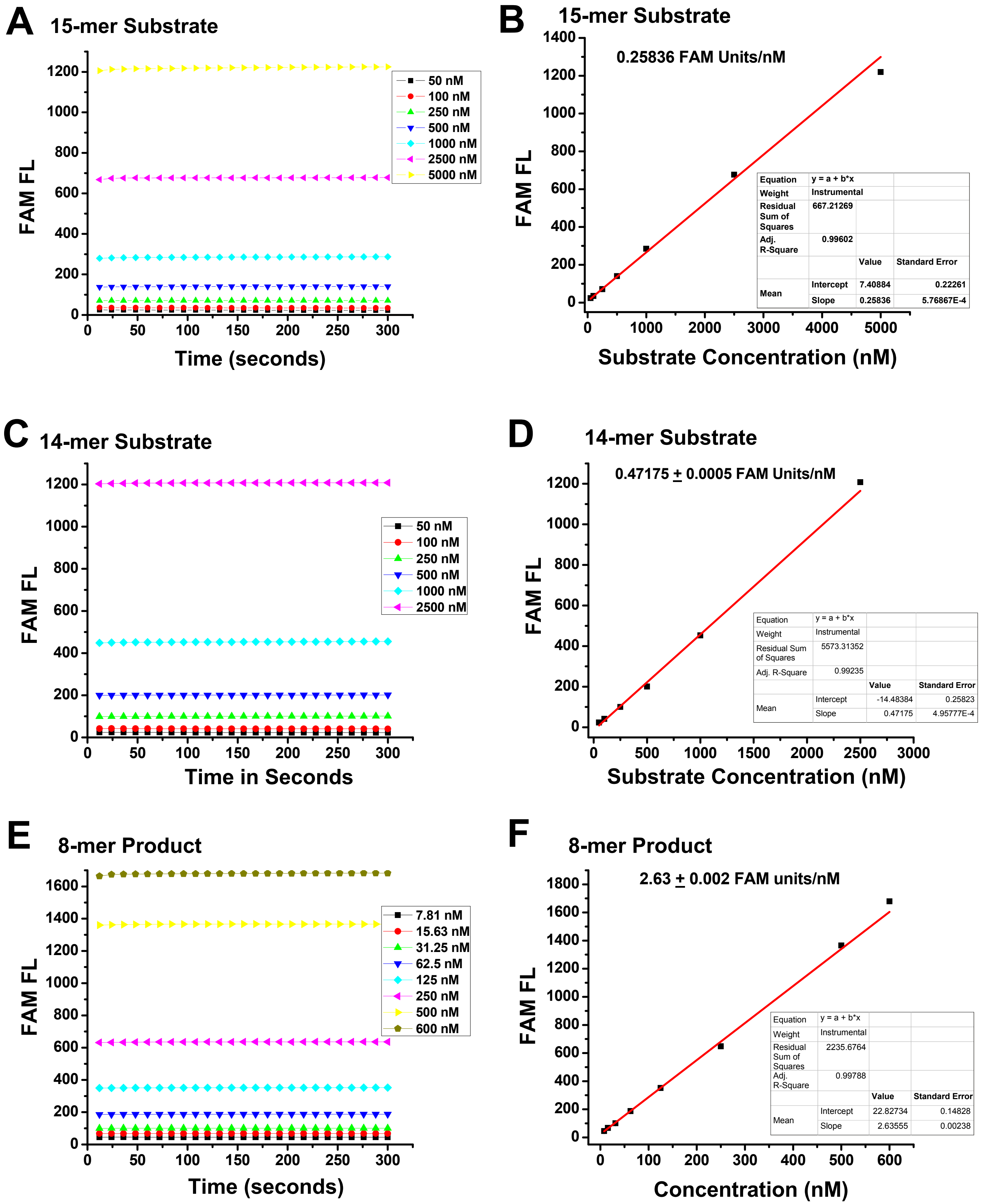

### Supplementary Figure 2, Myers and Sullivan

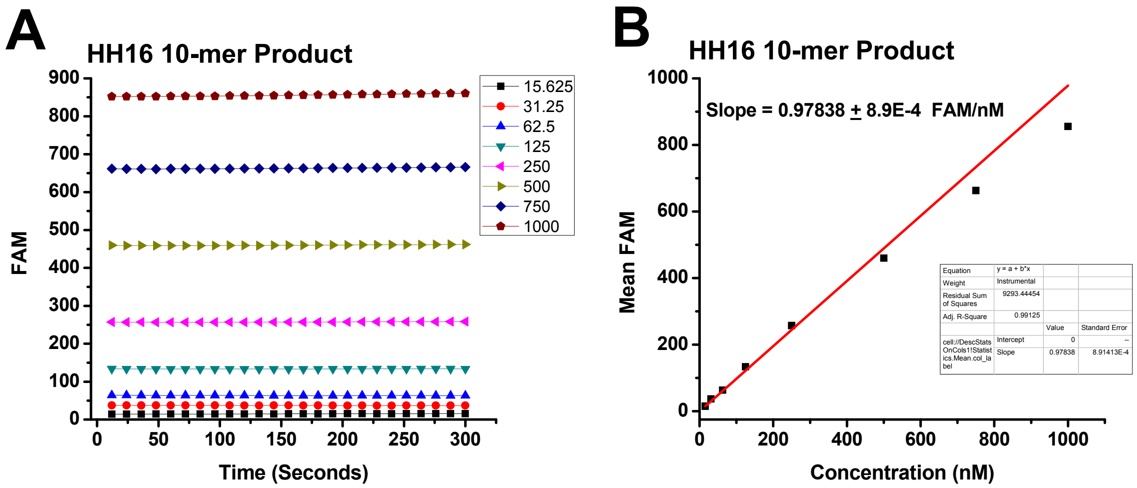

### Supplementary Fig 3, Myers and Sullivan

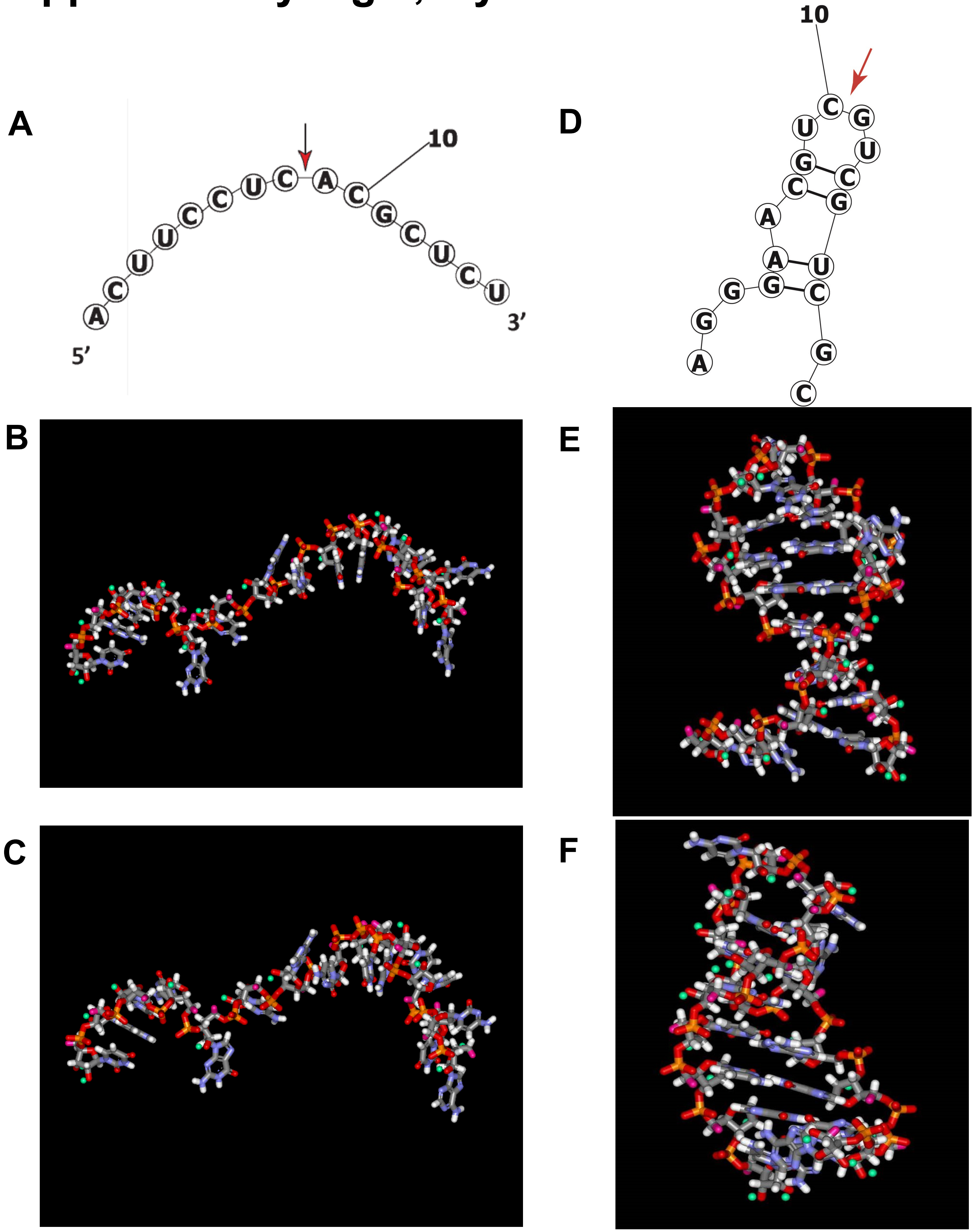

### Supplementary TABLE 1 Myers and Sullivan

#### HhRz Kinetic Model Parameters (Stage-Zimmermann and Uhlenbeck Model)

| Construct | $K_{d,1}$ | $k_{-1}$ | $K_{d,3}$ | $k_3$ | $K_{d,4}$ | $k_4$ | $K_{d,5}$ | $k_5$ | $K_{d,6}$ | $k_6$ |
|---|---|---|---|---|---|---|---|---|---|---|
| **WT (GAAA) 15-mer substrate** | 0.0021 | 1.03E-4 | 210 | 10.6 | 89.8 | 4.49 | 20.9 | 1.04 | 917 | 45.8 |
| **WT (GAAA) 14-mer substrate** | 0.0178 | 8.98E-4 | 212 | 10.6 | 89.8 | 4.49 | 294 | 14.69 | 917 | 45.8 |
| **A7U (GAAA/AGUA) 15-mer substrate** | 0.0043 | 2.15E-4 | 245.7 | 12.3 | 223 | 11.13 | 410 | 20.32 | 1060 | 53.13 |
| **A7U (GAAA/AGUA) 14-mer substrate** | 0.02754 | 1.38E-3 | 245.7 | 12.3 | 1679 | 83.95 | 387 | 19.35 | 1060 | 53.13 |

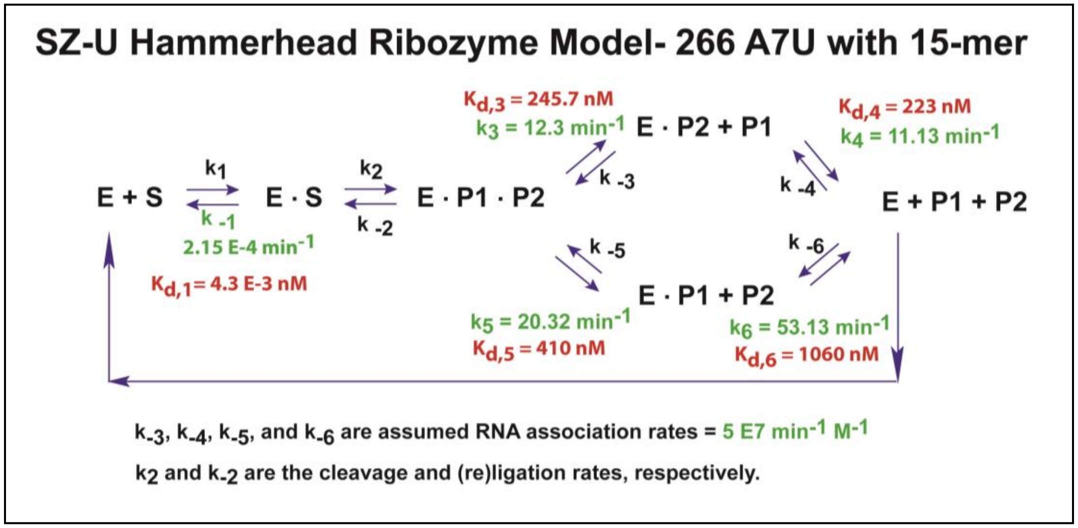


Table shows kinetic parameters calculated for Stage-Zimmermann and Uhlenbeck (1998) HhRz Model with Outcomes for the WT(GAAA) and A7U (GAAA/AGUA) *hRHO* 266 EhhRzs for both the 15-mer and 14-mer substrates. The kinetic model diagram (box) shows the Kd values and rate constants calculated for the A7U(GAAA/AGUA) EhhRzs solely for the 15-mer substrate. The A7U(GAAA) and A7U(AGUA) EhhRzs have identical antisense flanks and so have the same calculated kinetic properties. $K_d$ values have units of **nM**. Rate constants (***k***) have units of **min$^{-1}$**.
